# Ionic waves drive fluid flow by inducing spatiotemporal porosity variations in the brain interstitial space

**DOI:** 10.64898/2026.11.18.766825

**Authors:** Drik Sarkar, Saikat Mukherjee

## Abstract

We develop a model of interstitial fluid (ISF) dynamics in the brain driven by traveling ionic waves. The cortex is modeled as a 2D porous sheet with an absorbent matrix. A prescribed 10% sinusoidal porosity variation induces mean flow of *O*(1) *µ*m/s, even without an imposed pressure gradient. Including perivascular spaces (PVSs) increases the velocity and pressure gradient, and yields volumetric flux through arterial and venous PVSs, with maximum magnitude of 2 × 10^−2^ *µ*L min^−1^ mm^−1^. We show analytically, in the large-wavelength limit, that the porosity variation enters the pressure field as first- and second-order corrections. We then simulate a physiologically realistic spreading depolarization (SD) wave, finding that the induced porosity variation raises the mean ISF pressure to 4 mmHg, which falls to −0.7 mmHg during recovery. SD yields a peak velocity of *O*(10^−2^) *µ*m/s and a peak flux of 3 × 10^−3^ *µ*L min^−1^ mm^−1^ , and transiently stalls the PVS-driven flow. Although the SD wave produces a larger porosity change than the prescribed wave, it evolves slowly and drives weaker flow.

## 1 Introduction

Fluid flow in a porous medium with a spatiotemporally varying internal structure is important in many natural, biological, and engineered systems. Examples include soil, fibrous media, and biological tissues [1, 2, 3, 4, 5, 6]. The brain interstitial space exemplifies this class, with a solid phase of neuronal and glial cells, and an extracellular space filled with interstitial fluid (ISF) forming the fluid phase. The porosity, or volume fraction of this interstitial space has been measured to be *α* ≈ 0.2 [7], though it can vary dynamically across phases of sleep and wakefulness and with ionic activity in the brain tissue [8, 9, 10, 11, 12]. The connection between ISF flow and the dynamic behavior of tissue porosity is particularly important in the context of the “glymphatic system,” in which cerebrospinal fluid (CSF) flows through a network of perivascular spaces (PVSs), and is hypothesized to mix with the ISF to flush out metabolic waste products from the interstitial space [13, 14, 15, 16, 17].

Diffusion in the brain interstitium has long been thought to be the primary mechanism of solute and metabolic waste transport, and has been the subject of thorough investigations [7, 18]. Recent studies, however, indicate the possibility of bulk flow in the interstitium that transports solute at a rate comparable to, or even exceeding, that of diffusion [19]. Indeed, the buildup of these waste solute molecules has been correlated with neurodegenerative disorders like Alzheimer’s disease [20, 21, 22], implying that an alteration in the transport physics may be causatively linked to the progression of these disorders. The glymphatic hypothesis requires a pressure gradient imposed on the interstitium that can drive bulk ISF flow [23, 21, 22, 24]. Such flow is likely slow, pulsatile, and viscous, and has proven challenging to measure directly in experiments [25]. Indeed, a key existing bottleneck is experimentally resolving the ISF flow pathways, whose characteristic scales range from micrometers down to nanometers [26, 7, 23] and remain difficult to resolve even with two-photon microscopy, otherwise the gold standard for in vivo imaging [25, 27, 28].

In this paper we explore whether spatiotemporal variations in porosity due to ionic activity in the brain interstitium can provide an alternative mechanism to facilitate ISF flow and CSF-ISF exchange, in addition to the pressure-induced bulk flow. Such variations arise from ionic activities of the cells forming the solid matrix of the brain tissue. Ionic fluxes in the brain under physiological conditions create traveling chemical waves of varying wavelength, frequency, and amplitude, during different phases of sleep and wakefulness (delta, theta, alpha, gamma, and beta waves) [29, 30, 31, 32]. These traveling waves can shrink and expand the extracellular space by inducing cell swelling through osmotic gradients across the cell membrane that develop due to fluctuation in ions such as sodium, calcium, and potassium [11, 33, 34, 35, 10, 36, 37]. Such waves also arise during stroke, traumatic brain injury, migraine, headache, seizures, cardiac arrest, and hemorrhage [38, 39, 40, 41, 42, 43]. The swelling is driven by influx of ions like Na^+^ and Ca^2+^ into the cells and an efflux of K^+^ [33, 34, 35, 10]. This ionic redistribution across the cell membrane leads to osmotic water influx into the cell from the extracellular space. Additionally, the excess K^+^ in the cortex may lead to changes in the lumen radius of the arteries, boosting perivascular CSF flow [42]. In acute conditions, the excess ions in the cortex then spread by diffusion along the cortical surface, leaving behind a trail of swollen, depolarized neurons and constricted vasculature [40]. The resulting spreading depolarization (SD) wave has a typical speed of *O*(1) mm/min [44, 33, 45, 42, 25].

Recent studies provide evidence of traveling-wave-induced alterations in CSF and ISF dynamics under both physiological and acute conditions. A recent biophysical model reveals electrochemical gradients induced by neuronal activities can generate fluid velocities of typical magnitudes of 14 *µ*m/min in the extracellular space [46]. Recent experiments further provide evidence that neuronal ionic activity and externally applied neural stimulation can drive ISF flow [47, 48]. A recent murine experiment demonstrated that spreading depolarization waves induced by ischemic stroke in mice result in large CSF influx into the cortex, a novel discovery indicating that CSF is the earliest contributor to brain edema (water accumulation in tissue) following stroke [40]. Using CSF tracers and two-photon microscopy, another study found fluctuations in CSF and ISF volumes over the course of seizure progression [43], specifically decreasing post-seizure, which increased tau protein aggregation. Vasoconstriction has been demonstrated as a primary mechanism driving perivascular CSF flow following depolarization and has since been investigated numerically [25, 42]. However, cell swelling is an additional characteristic feature that can drive ISF flow [49, 50, 51], which coupled with enhanced perivascular flow may contribute jointly to edema and blood-brain-barrier compromise under acute conditions.

Spatiotemporal porosity variations as a flow-inducing mechanism has been studied in several contexts, particularly in geophysical settings and fibrous media [52, 4, 5]. In many problems such as magma migration, an evolving “porosity wave” is produced by localized fluid production or absorption [52, 53], which enters as a source term in the continuity equation. Another closely related scenario is fluid flow in unsaturated absorbent porous materials, such as fiber bundles and paper towels, where a “wicking” front forms as liquid is absorbed by the fibers [5, 4]. Overall, these cases can be categorized as “dual-scale” porous media, where the fluid dynamics is governed by two length scales: the pore-length scale *L*, the distance between pores, and the imbibition length scale *l* inside the solid matrix, as shown by the schematic in Fig. 1.

**Figure 1:**
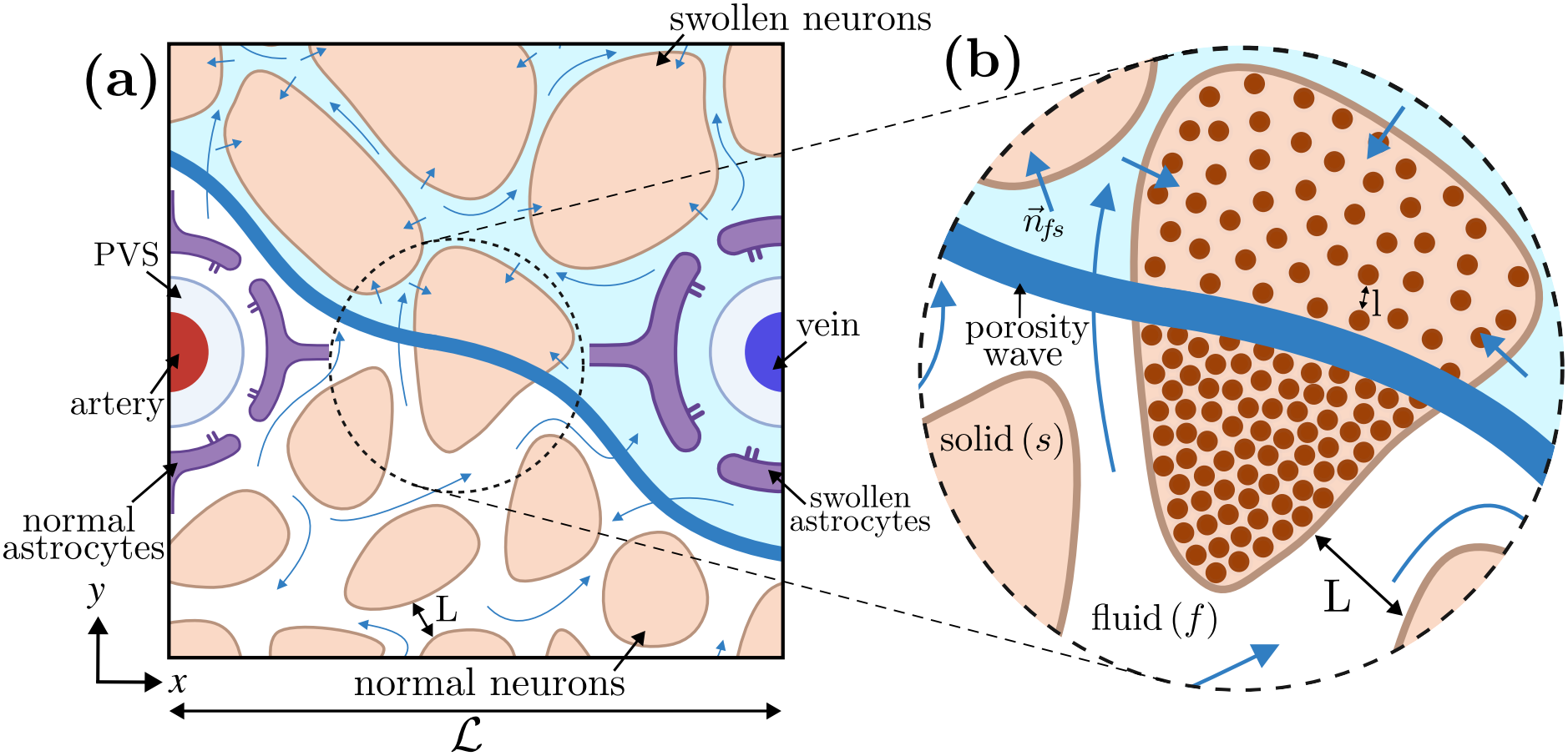
Computational and volume-averaging domains. (a) A 2D square region of the brain cortex of length ℒ = 250 *µ*m, showing neuronal cells, glial cells, and PVSs. An ionic wave (blue) separates swollen from non-swollen cells. (b) Representative elementary volume (REV) of the domain used for volume averaging, showing fluid and solid phases.

In this paper we show how the dual-scale porous media framework can be implemented to model ISF flow subjected to traveling ionic waves in a domain representing the brain tissue. The wave swells the cells on which it is incident, locally decreasing the porosity, and leaves a trail of swollen cells in its wake, with higher porosity in the tissue ahead of it. Once the wave has passed, the cells shrink and the porosity recovers to its baseline value. We study a theoretically prescribed porosity variation and porosity variation obtained from a physiologically realistic model of spreading depolarization [42]. The analytical framework developed here is generalized and can be readily extended to any porous medium undergoing arbitrary spatiotemporal porosity fluctuations. We first describe our approach in Sec. 2, which includes a discussion of the simulation domain and governing equations. We then describe our results in Sec. 3, starting with the fluid dynamics induced by a prescribed porosity variation and then by a simulated spreading depolarization wave. Lastly, we present our concluding remarks in Sec. 4.

## 2 Approach

Figure 1 illustrates the physical system and computational domain under consideration. The computational domain, denoted Ω, is a 2D square sheet of brain interstitium between a penetrating arteriole and venule, as shown in Figs. 1(a) (*x*-*y*-plane). We take this domain as a portion of an infinite tiling of arterioles and venules, stacked along the cortical depth (*z*-axis) [22, 21, 24]. We also evaluate cases where PVSs are removed. The characteristic length is the typical arteriole-to-venule distance in primates, *ℒ* = 250 *µ*m [54, 55, 22, 21]. The schematic shows an ionic traveling wave in blue, separating the lower porosity region with swollen cells and the region with non-swollen cells. Figure 1(b) shows the representative volume (REV) of arbitrary radius *r*_0_ and volume 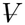, used for volume averaging. The three length scales under consideration are the macroscopic length scale, *ℒ*, the pore-length scale, *L*, and the absorption or imbibition length scale, *l*, such that *l < L < ℒ*. The REV radius satisfies *l < L* ≤ *r*_0_ ≤ *ℒ* [56, 57]. *A*_*fs*_ is the surface area of the fluid-solid interface and 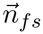 is the normal directed from the fluid to the solid phase.

### 2.1 Governing equations of fluid flow

We next describe the governing equations of the macroscopic fluid flow in the interstitium. A detailed treatment of volume averaging of the mass and momentum equations is provided in the Appendix A, and here we present the reduced equations. The pore-scale fluid velocity is 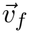 , while the local variation of the solid and fluid phase is captured by the velocity of the fluid-solid interface 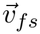. Volume averaging the continuity equation for incompressible, single-phase flow in a porous medium yields,

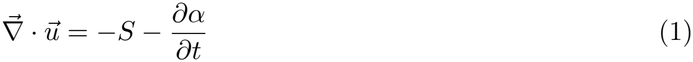

where 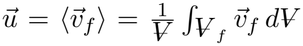 is the volume-averaged fluid velocity, 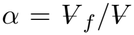 is the porosity, and 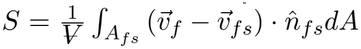 is the imbibition rate acting as a sink term related to the ISF imbibition by the cells. During swelling, *∂*_*t*_*α <* 0, acting as a positive source term to Eq. (19). This is because when interstitial volume decreases, by volume conservation, the fluid flux through the constrictions increases. An opposite behavior is expected during de-swelling. Assuming the *S* is linearly proportional to the rate of change of the solid volume fraction, as typically employed in paper and fiber matrices [5, 4], 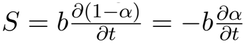 , where 0 ≤ *b* ≤ 1 is the absorption coefficient, and the mass conservation equation Eq. (19) reduces to,

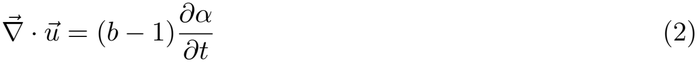

Equation (2) implies that when *b* = 1, or when the solid matrix expands at the same rate as absorption, a divergence-free volume-averaged flow is maintained. However if *b <* 1, the storage effect of the solid matrix to the volume-averaged flow is nontrivial. We have used *b* = 0.9 following Ref. [5] on fibrous porous media. Volume averaging the conservative form of the Navier-Stokes equation in the fluid phase, neglecting any body forces, yields a modified Darcy’s law [56],

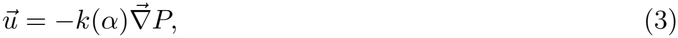

where *k*(*α*) = *K*(*α*)*/µ* is the hydraulic conductivity of the brain tissue, *µ* is the coefficient of dynamic viscosity, and *P* = ⟨*p*⟩^*f*^ is the intrinsic average of the pressure. This reduction leverages the negligible Darcy number 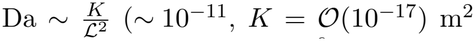 [23] and *ℒ* = *O*(10^−3^) m) and negligible Reynolds number, Re = |*U* |*L/v* ∼ *O*(10^−6^) (*U* ∼ 1*µ*m*/*s, and *v* = 10^−6^m^2^*/*s). By combining the mass (Eq. 2) and the momentum conservation equations (Eq. (3)), we obtain a Poisson–type equation for the pressure field

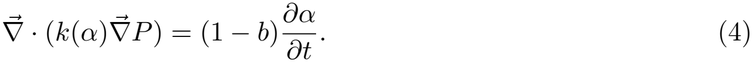

Equation (4) indicates that the flow can be driven even without an externally imposed pressure difference, and solely by spatiotemporal variations in the porosity and permeability. We use a Kozeny–Carman-type relation [58] to relate the permeability and porosity, which has been used in the context of ISF flow in the brain cortex previously [21, 12]

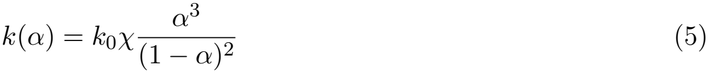

where 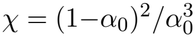, with *α*_*0*_ *= 0*.*2* as the reference porosity [7], and *k*_0_ = 2*×*10^−6^ cm^2^mmHg^−1^s^−1^ [24] is the corresponding baseline hydraulic conductivity.

### 2.2 Spatiotemporal variations in porosity

We investigate two types of porosity variation. The first one is a prescribed variation of the form,

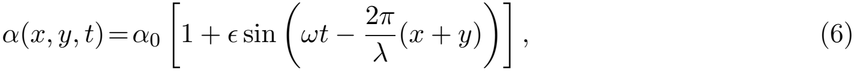

where *ω* = 2*π f* is the angular frequency and *λ* is the wavelength. Equation (6) yields a traveling porosity wave that enters the domain at the top-right corner ((*x, y*) = (*ℒ, ℒ*)), and exits at the bottom left-corner ((*x, y*) = (0, 0)). This models porosity fluctuations expected by physiological brain waves, originating in different phases of sleep and wakefulness, namely alpha, beta, theta, and gamma waves [29, 30, 31, 32, 47]. Here, the intracellular space is considered as the solid phase 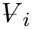, the extracellular space as the fluid phase 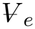, and the porosity is defined as 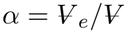, where 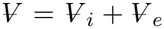. We choose *λ* = 1 mm and frequency *f* = 1 Hz, corresponding to a delta wave in mice [30, 31, 59, 32], and an amplitude of *ϵ* = 0.1, or a 10% variation from the reference value of *α* _0_ = 0.2.

We also study porosity variation induced by a spreading depolarization (SD) wave. Acute depolarization waves during stroke, migraine, seizures, and anoxia can be modeled as a multi-species nonlinear system of ionic reaction-diffusion equations [11, 10, 60, 33, 44]. In a recent study, we have modeled SD [42] using physiologically realistic ionic dynamics of potassium and sodium ions, by solving Hodgkin–Huxley and Goldman–Hodgkin–Katz kinetics, with membrane gating protein dynamics, in the intracellular space and ion diffusion in the extracellular space [42]. Here, we extend this to 2D and incorporate porosity variations induced by osmotic cell swelling. The full reaction-diffusion system is presented in Eq. (67)-(72) (Appendix D). The porosity variation induced by ionic redistribution across the cell membrane is:

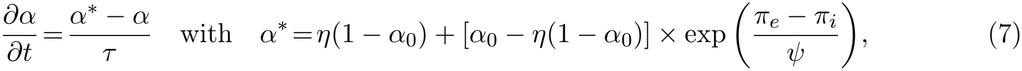

where *π*_*i*_ and *π*_*e*_ are the intra- and extracellular osmolarities given as,

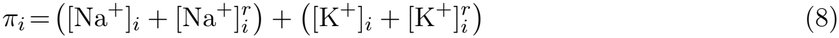

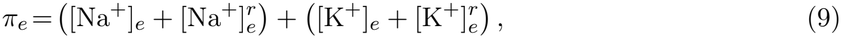

where the superscripts *r* denotes the resting values, provided in Appendix D.

## 3 Results and discussions

### 3.1 Prescribed porosity variation

We begin the results by investigating the prescribed porosity wave Eq. (6). Figure 2(a) shows the porosity field at four temporal phases, *ωt* = 0, *π*/2, *π* and 3*π*/2. The corresponding spatiotemporal pressure fluctuations are shown in Fig. 2(b). The porosity field induces an instantaneous pressure gradient along the diagonal of the domain that drives ISF flow. Because of the no-flux material walls, we fix the gauge pressure using ⟨*P*⟩_Ω_ = 0 for all times, as shown by the color contours. However, the pressure gradient is nonzero and is driven by the porosity variation. The pressure gradient is steep at *ωt* = 0, even though the mean porosity is lower. It then reduces and remains approximately constant for *ωt* = [*π*/2, *π*, 3*π*/2]. This dependence follows Eq. (4), which implies that the pressure gradient magnitude scales as 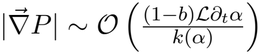, where ℒ is the length scale and 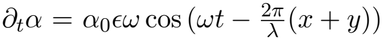. Since ∂_*t*_*α* is large at *ωt* = 0 while *α* and consequently *k*(*α*) are small, the pressure gradient peaks at this phase. Additionally, the reciprocal dependence on *k*(*α*) implies that when porosity and the hydraulic conductivity increase, the pressure gradient drops at *ωt* = [*π*/2, 3*π*/2].

**Figure 2:**
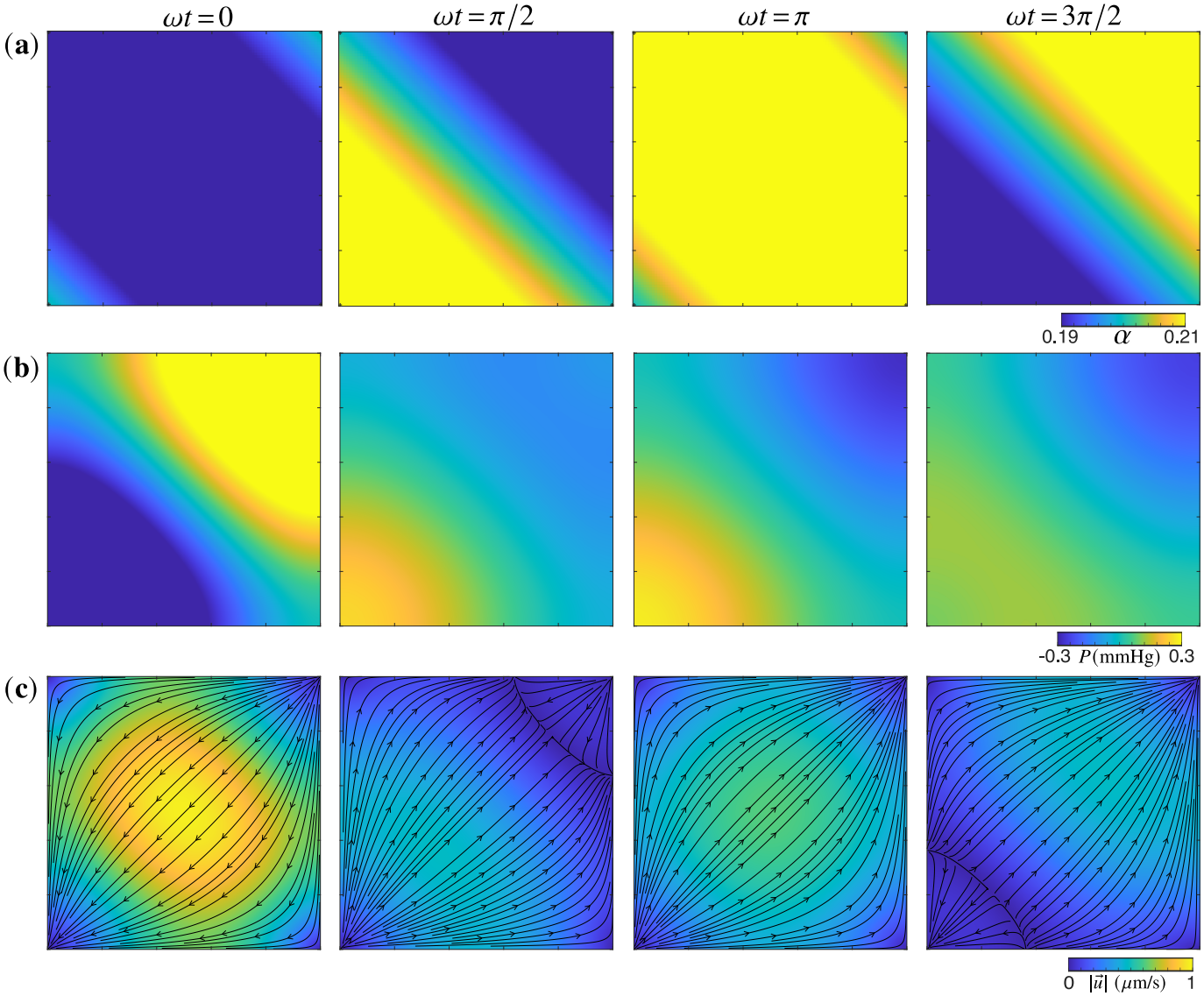
Fluid flow driven by the prescribed porosity variation Eq. (6) in a domain without PVSs at phases *ωt* = 0, *ωt* = *π*/2, *ωt* = *π*, and *ωt* = 3*π*/2. Color contours of (a) porosity field, (b) pressure field, and (c) velocity magnitude with streamlines overlaid.

The induced flow field is shown in Fig. 2 (c) and is driven by the pressure gradient from top-right to the bottom-left of the domain. The large pressure gradient at *ωt* = 0 drives substantial fluid flow of magnitude exceeding 1 *μ*m/s at the center of the domain. The flow weakens and starts reversing at *ωt* = *π*/2. The direction of the flow is fully reversed at *ωt* = *π*, relative to *ωt* = 0. Although the pressure gradient is comparatively small at *ωt* = *π*, when compared to *ωt* = 0, the velocity magnitude is substantial, driven by the high permeability. A second transitional phase is at *ωt* = 3*π*/2 (direction reversed relative to *ωt* = *π*/2), before the cycle repeats.

We next study a prescribed porosity variation in a parenchymal section between an arterial and a venous PVS, held at a constant pressure difference Δ*P*_PVS_. Two cases are considered: Δ*P*_PVS_ = 0 mmHg (no pressure difference between the PVSs), and Δ*P*_PVS_ = 0.4 mmHg. Prior numerical studies have used Δ*P*PVS = *O*(1) mmHg to investigate bulk flow in the interstitium [21, 22, 24, 23]. A baseline case of ISF flow driven solely by a PVS pressure difference Δ*P*_PVS_ = 0.4 mmHg, is shown in Fig. 14 for reference. The color contours of the pressure field are shown in Fig. 3(a) and (b). In both the cases, the porosity-induced pressure superposes on the existing field, yielding spatially nonuniform pressure variation at *ωt* = 0 and *ωt* = *π*, with the gradient reversed in the latter. At *ωt = π/2*, the pressure inside the domain falls below the fixed boundaries, and becomes negative, so the pressure gradient points inward. This reverses at *ωt* = 3*π*/2, when the domain pressure exceeds the boundaries. When Δ*P*_PVS_ = 0.4 mmHg, the pressure gradient is steeper, at all times, than Δ*P*_PVS_ = 0 mmHg, as expected.

**Figure 3:**
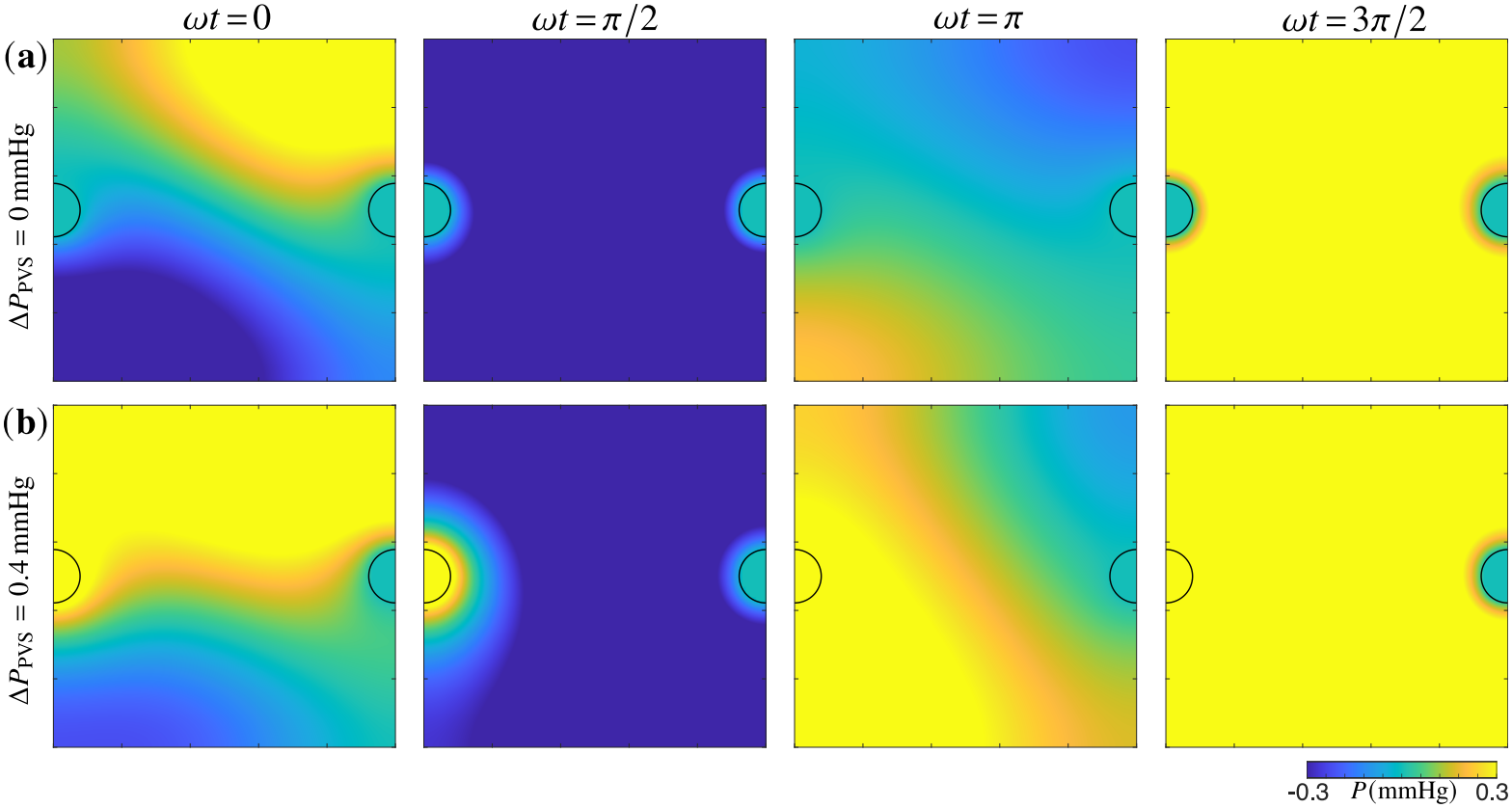
Color contours of the pressure field induced by the prescribed porosity variation Eq. (6) in a domain with PVSs at phases *ωt* = 0, *ωt* = *π*/2, *ωt* = *π*, and *ωt* = 3*π*/2. (a) Δ*P*_PVS_ = 0 mmHg, and (b) Δ*P*_PVS_ = 0.4 mmHg.

This pressure field drives ISF flow in and out of the domain through both PVSs as shown in Fig. 4(a) and (b). At *ωt* = 0, flow is drawn in from the arterial PVS and is driven out from the venous PVS due to the nonuniform pressure gradient. As the phase advances to *ωt* = *π*/2, the reduced domain pressure draws fluid inward from both the PVSs. Interestingly, as the wave travels diagonally across the domain, it draws more from the arterial than the venous PVS. At *ωt* = *π*, the pressure distribution and the flow direction are inverted relative to that at *ωt* = 0. At *ωt* = 3*π*/2, the domain pressure exceeds the PVSs, driving ISF outward through both, with more efflux through the venous PVS because of the wave propagation direction. For Δ*P*_PVS_ = 0.4 mmHg, fluid is driven from the arterial to venous PVS at all times due to the perivascular pressure difference, with the porosity variation superposed on top. Consequently, the inflow through the arterial PVS is moderately faster for Δ*P*_PVS_ = 0 mmHg, while the outflow through the venous PVS is faster for Δ*P*_PVS_ = 0.4 mmHg, as seen by comparing Fig. 4(a) and (b).

**Figure 4:**
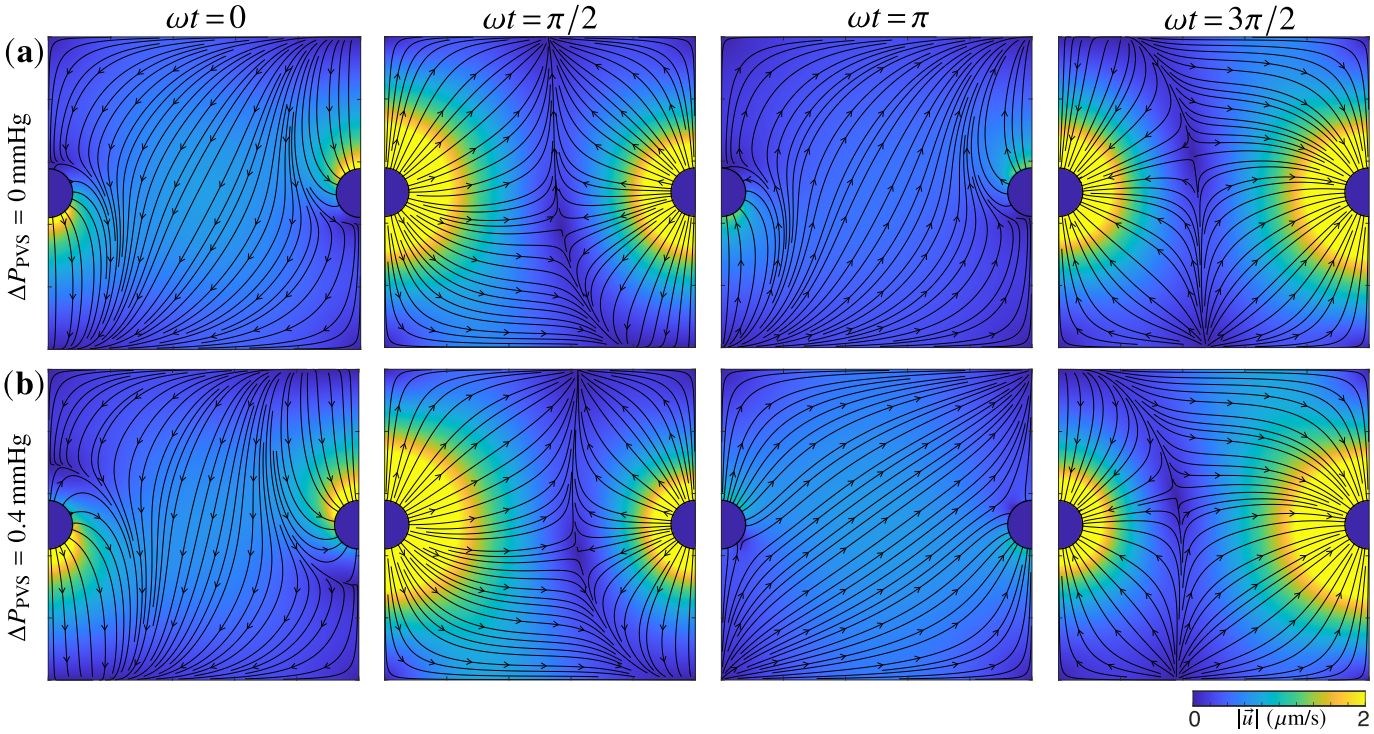
Color contours of the velocity magnitude with streamlines overlaid, induced by the prescribed porosity variation in a domain with PVSs at phases *ωt* = 0, *ωt* = *π*/2, *ωt* = *π*, and *ωt* = 3*π*/2. (a) Δ*P*_PVS_ = 0 mmHg, and (b) Δ*P*_PVS_ = 0.4 mmHg.

We plot the temporal variations of mean pressure, pressure gradient magnitude, and velocity magnitude in Fig. 5. Panel (a) shows the mean magnitude of the pressure gradient 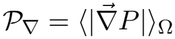 in the absence of PVSs, which reaches a peak of *P*_∇_ = 4.36 × 10^−3^ mmHg/μm at *ωt* = 0 when ∂_*t*_*α* is maximum and *k*(*α*) and *α* are minimum. It reaches a minimum *P*_∇_ ≈ 9.69 × 10^−4^ mmHg/μm near *ωt* ≈ [*π*/2, 3*π*/2], where ∂_*t*_*α* is minimal. The minimum value does not correspond exactly to *ωt* = [*π*/2, 3*π*/2], since the temporal phase is *θ* = *ωt* ™ (2*π*/*λ*)(*x* + *y*). *P*_∇_(*t*) then rises and reaches local maxima at *ωt* ≈ [3*π*/4, 5*π*/4], before the cycle repeats. The increase in the magnitude for *π*/2 ≲ *ωt* ≲ 3*π*/2 is related to the increase in *k*(*α*). The mean velocity magnitude follows an analogous trend and varies between 0.18 μm/s ≲ 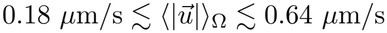 (Fig. 5(b)).

**Figure 5:**
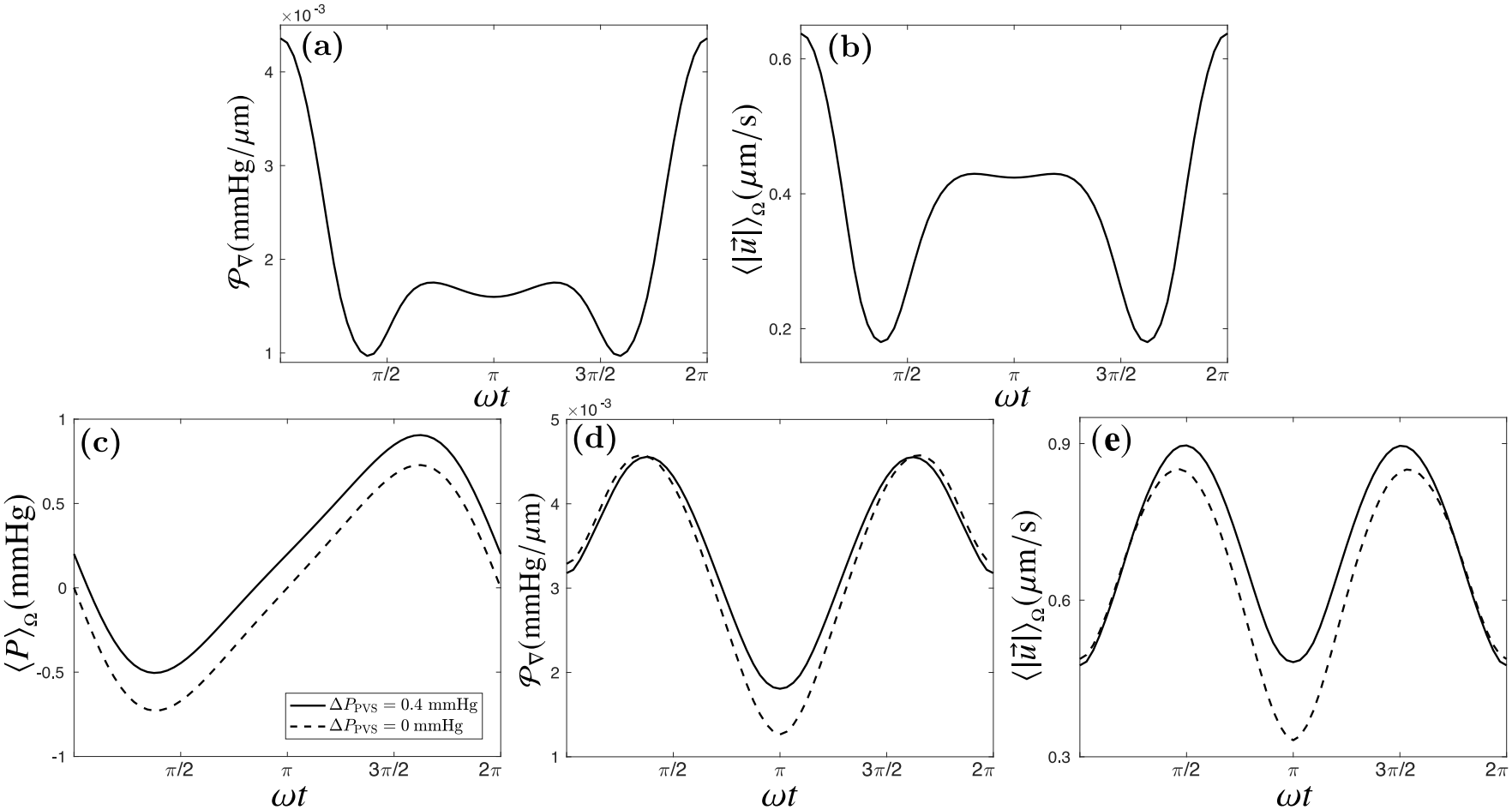
Temporal variation of pressure and velocity induced by a prescribed porosity variation Eq. (6). (a) Mean magnitude of pressure gradient *P*_∇_ and (b) mean velocity magnitude in a domain without PVSs. (c) Mean pressure, (d) *P*_∇_, and (e) mean velocity magnitude in a domain with PVSs. The solid and dotted curves correspond to Δ*P*_PVS_ = 0.4 mmHg and Δ*P*_PVS_ = 0 mmHg.

Figure 5(c-e) shows the temporal variations of ⟨*P*⟩_Ω_, *P*_∇_, and mean velocity magnitude 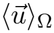, in the domain with PVSs. The mean pressure ⟨*P*⟩_Ω_ varies sinusoidally between −0.73 mmHg ≤ ⟨*P*⟩_Ω_ ≤ 0.73 mmHg for Δ*P*_PVS_ = 0 and between −0.50 mmHg ≤ ⟨*P*⟩_Ω_ ≤ 0.91 mmHg for Δ*P*_PVS_ = 0.4, with troughs and peaks near *ωt* = *π/*2 and *ωt* = 3*π/*2, as shown in Figure 5(c). The fluctuations are superimposed on the mean pressure due to the PVS boundaries and vanish, when averaged over a cycle. Indeed, at the leading order, the PVS pressure difference dictates the domain pressure, and the porosity-induced effect enters as first and second-order corrections in Δ*α*. We demonstrate this analytically in the large-wavelength limit in Appendix C.

The temporal variation of *P*_∇_ in the domain with PVSs is shown in Fig. 5(d) and shows peaks near *ωt* = [*π/*2, 3*π/*2]. The peaks correspond to phases in which the domain is at a smaller or larger pressure than the fixed boundaries. The troughs near *ωt* = [0, *π*, 2*π*], correspond to phases in which the domain pressure is comparable to the PVS boundaries. For 0 ≤ *ωt* ≲ *π/*2 and 3*π/*2 ≤ *ωt* ≲ 2*π, P*_∇_ is moderately larger forΔ*P*_PVS_ = 0 (dotted line), because of reduced pressure gradient around the arterial PVS at Δ*P*_PVS_ = 0.4 mmHg. Additionally, *P*_∇_ drops to a lower value for Δ*P*_PVS_ = 0 because Δ*P*_PVS_ *>* 0 sustains a pressure gradient even when the porosity-induced gradient is minimal. The mean velocity magnitude follows the same trend and is plotted in Fig. 5(e).

We next quantify the volumetric flux through the PVS boundaries, relevant to CSF–ISF mixing, as the surface integral of the fluid velocity dotted with the outward normal at the arterial PVSΓ_*a*_ and venous PVSΓ_*v*_,

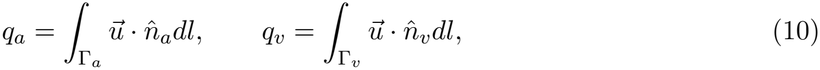

where, *q*_*a*_ or *q*_*v*_ corresponds to the volume flow rate per unit depth of the cortex leaving the arterial or venous PVS respectively. With the normal unit vector pointing outward, any influx yields *q <* 0 whereas any efflux yields *q >* 0. Figure 6 plots the volumetric flux through the arterial (solid red) and venous (solid blue) PVSs driven by the prescribed porosity variation for Δ*P*_PVS_ = 0.4 mmHg. For 0 ≤ *ωt* ≤ *π*, fluid enters the domain through both PVSs because the porosity wave lowers the domain pressure below the PVS boundaries. For 0 ≤ *ωt* ≤ *π*, the magnitude of influx through the arterial PVS is larger because of the diagonal wave travel direction. For *π* ≤ *ωt* ≤ 2*π*, fluid exits the domain through both the PVSs, with greater efflux through the venous PVS. The corresponding results for Δ*P*_PVS_ = 0 mmHg (dashed curves) indicate that both the venous efflux and the arterial influx are smaller than Δ*P*_PVS_ = 0.4 mmHg. For reference, the dashed-dotted lines indicate the fluxes through the PVSs driven by Δ*P*_PVS_ = 0.4 mmHg without any porosity variation. In the presence of porosity variation, for Δ*P*_PVS_ = 0.4 mmHg, the maximum arterial influx is |*q*_*a*_|_*max*_ = 1.84 *×* 10^−2^ *µ*L min^−1^ mm^−1^ and the maximum venous efflux is |*q*_*v*_|_*max*_ = 1.62 *×* 10^−2^ *µ*L min^−1^ mm^−1^.

**Figure 6:**
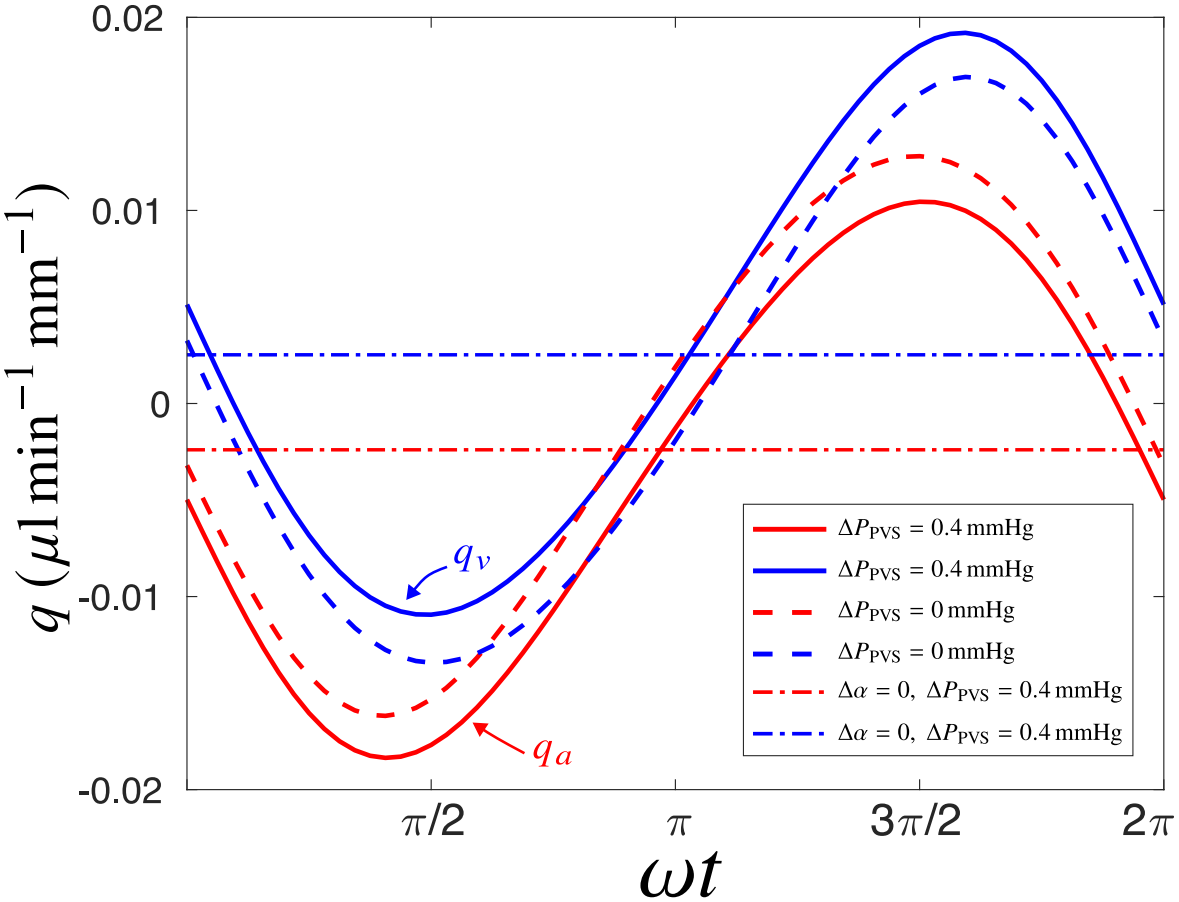
Temporal variation of the volumetric flux through the PVSs driven by prescribed porosity variation Eq. (6).

### 3.2 Porosity variation induced by a simulated spreading depolarization wave

Cortical spreading depolarization (SD) begins when ion regulation breaks down in a small cortical region, causing cells to depolarize almost completely [41]. The disturbance does not remain localized. It moves slowly through the gray matter, typically at velocities of 2–6 mm/min [42, 33, 44], as extracellular potassium released from one region diffuses and triggers depolarization in the next. This potassium front produces localized osmotic gradients that induce cell swelling driving porosity variations and ISF flow. In our model, the SD wave is initialized near the upper-right corner of the domain using a line-shaped Gaussian profile, given by Eq. (27), and then evolves according to the reaction-diffusion system Eq. (67)-(72).

Figure 7(a) and (b) show the spatiotemporal evolution of the extracellular potassium and sodium concentrations, [K^+^]_*e*_ and [Na^+^]_*e*_. SD results in a fast wave that propagates diagonally into the domain, and within approximately 3 s, the potassium concentration in the whole domain rises to 50 mM from the resting value 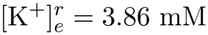, while the extracellular sodium concentration falls to 80 mM from 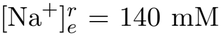. We define the start of the recovery phase to be *t*_*SD*_ = 41.9 s, when K^+^]_*e*_ goes back to the resting value. Beyond *t*_*SD*_, [K^+^]_*e*_ drops moderately below 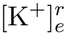. We terminate the simulations at *t* = 100s, when the membrane potential is within 12% of its resting value of -70 mV. We denote *t*_*SD*_ ≤ *t* ≤ 100 s as the recovery time, although the recovery remains ongoing beyond *t* = 100 s, which we do not investigate further.

**Figure 7:**
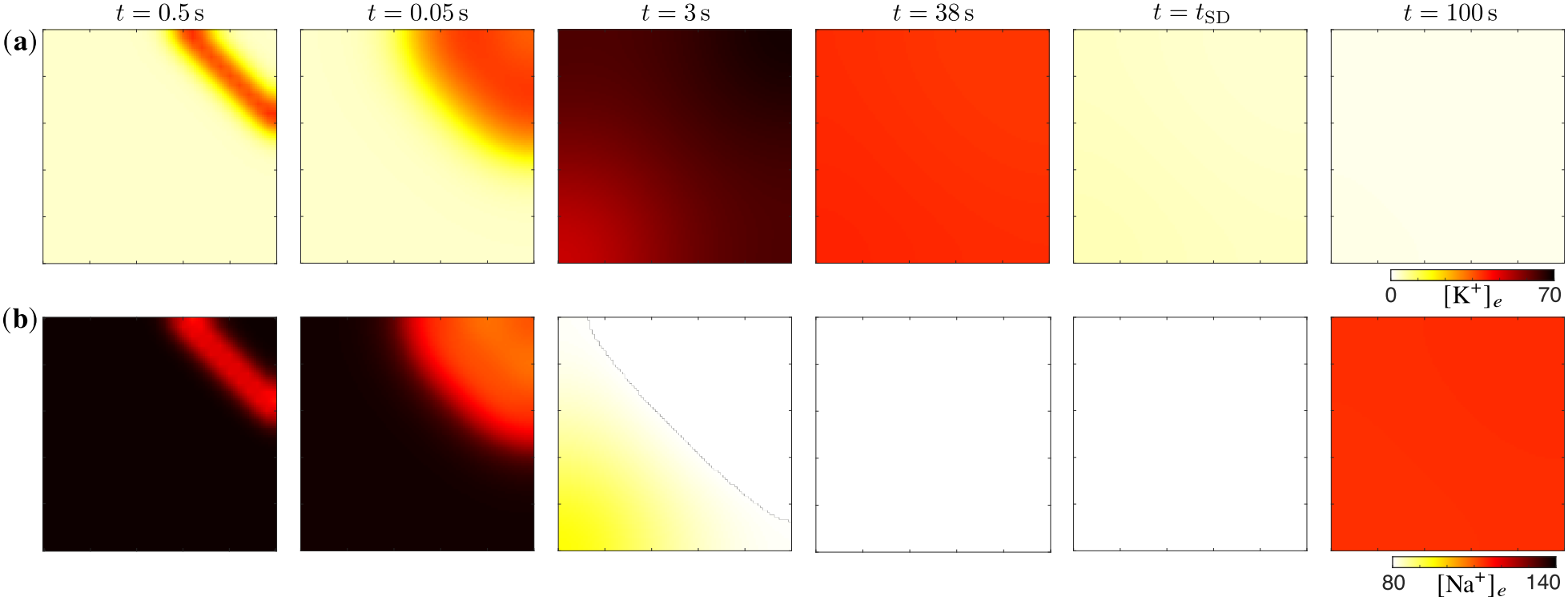
Ion concentration fields produced by a simulated spreading depolarization wave propagating through the domain. Color contours of (a) extracellular potassium ion concentration ([K^+^]_*e*_) and (b) extracellular sodium ion concentration ([Na^+^]_*e*_) are shown at five different time instances during the propagation of the depolarization wave, *t* = 0.05 s, 0.5 s, 3 s, *t*_*SD*_ = 41.9 s, and 100 s going from left to right.

Figure 8(a) shows the color contours of the porosity field induced by the SD wave at the same time instances as Fig. 7. The initial potassium spike induces a localized, transient increase of *α*, indicated by bright yellow at *t* = 0.05 s and *t* = 0.5 s in Fig. 8(a). As the potassium front invades the domain further, porosity drops. The mean porosity falls to a minimum of 0.03 at *t*=*t*_*SD*_, beyond which it increases again (Figure 8(b)). Figure 8(c) shows the spatially averaged rate of porosity change, which is negative as the tissue swells. It increases in magnitude at early time and stays constant at ≈ −4.5 *×* 10^−3^ s^−1^ for *t* ≲ 25 s, beyond which it weakens to 0 at *t*_*SD*_. For *t* ≳ *t*_*SD*_, the value becomes positive ∼ 10^−3^ s^−1^, which stays constant through the end of the simulation.

**Figure 8:**
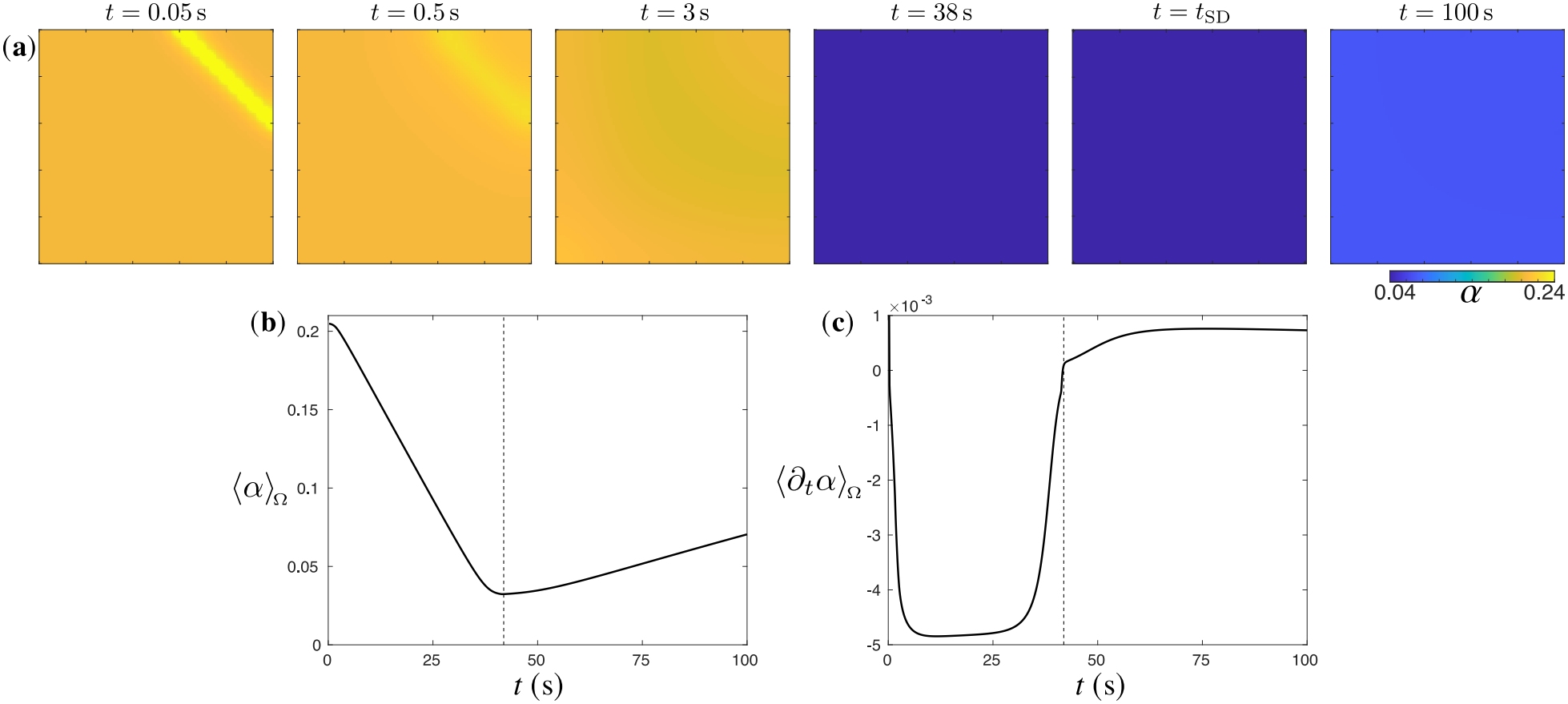
Porosity variation due to a spreading depolarization wave. (a) Color contours of porosity field *α*(*x, y, t*). Times correspond to the instances shown in Fig. 7. (b) Mean porosity as a function of time, and (c) mean rate of porosity change versus time.

Figure 9(a) shows the pressure induced by the SD-induced porosity in a domain without PVSs.

**Figure 9:**
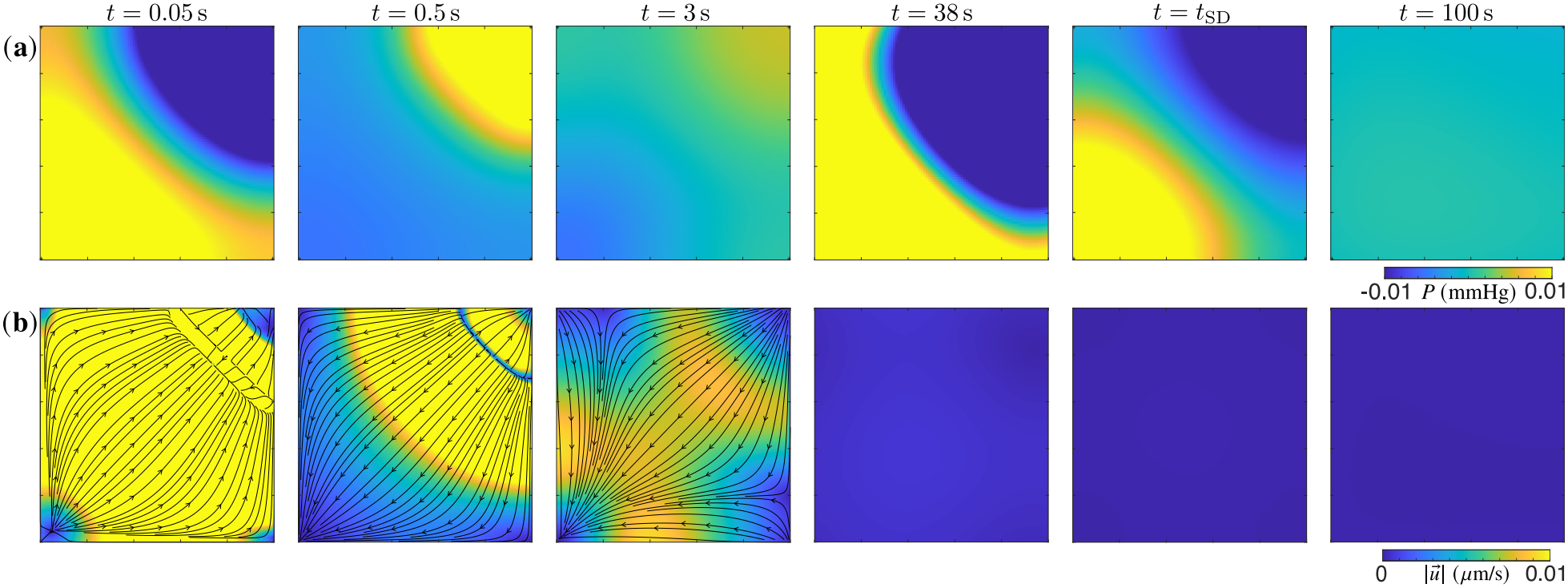
Pressure variation and fluid flow driven by a spreading depolarization wave. Color contours of (a) pressure field in a domain without PVSs, and (b) corresponding velocity magnitude with streamlines overlaid. Times correspond to the instances shown in Fig. 7.

At the start, pressure drops at the wave-initiation site (blue in the *t* = 0.05 s panel) with the pressure gradient pointing from bottom-left to top-right of the domain. The pressure peaks at *±*0.01 mmHg, with the gauge fixed such that mean pressure is zero. The pressure gradient then turns abruptly, pointing from top-right to bottom-left of the domain, aligning with the SD-wave propagation direction for *t < t*_*SD*_. As the potassium front exits the domain, the direction of the pressure gradient recovers back (*t* = 38 s). As 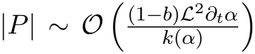 , localized regions of high porosity (and conductivity *k*(*α*)) reduce the pressure value, while pressure is largest during the swelling phase when |*∂*_*t*_*α*| is greatest, and decreases thereafter. The resulting ISF flow is shown in Fig. 9(b), using color contours of the velocity magnitude with streamlines overlaid. At the start, the fluid rushes towards the SD initiation site, after which the flow reverses, aligning with the direction of SD-wave travel. The velocity magnitude weakens over time, and vanishes for *t* ≳ *t*_*SD*_, reaching a peak 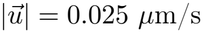 during wave-initiation.

The induced pressure field in the domain with PVSs, for Δ*P*_PVS_ = 0 and Δ*P*_PVS_ = 0.4 mmHg, are shown in Fig. 10(a)–(b). Since the SD-induced pressure magnitude is *P* ∼ *O*(10^−2^) mmHg, and it enters the dynamics as first order in Δ*α*, its contribution appears moderate at early times when the color range is −0.3 mmHg ≤ *P* ≤ 0.3 mmHg. For Δ*P*_PVS_ = 0, when the SD wave is initiated, the domain pressure becomes negative, resulting in an inward pointing pressure gradient from the boundaries (*t* = 0.05 s, in Fig. 10(a)). As time progresses, the domain pressure matches the PVS boundaries (*t* = 0.5 s and *t* = 3 s), then increases in magnitude to exceed them (*t* = 38 s). As the potassium front exits the domain, the domain pressure falls to negative values and result in weak inward-pointing pressure gradients that persist through the course of the simulation. For Δ*P*_PVS_ = 0.4 mmHg, the arterial PVS produces a larger gradient at initiation, after which the pressure field becomes nearly uniform, and increases with time (*t* = 0.5 s, *t* = 3 s, and *t* = 38 s, in Fig. 10(b)). For *t* ≳ *t*_*SD*_, the pressure in the domain drops again and the bulk PVS pressure driven flow is restored at long times.

**Figure 10:**
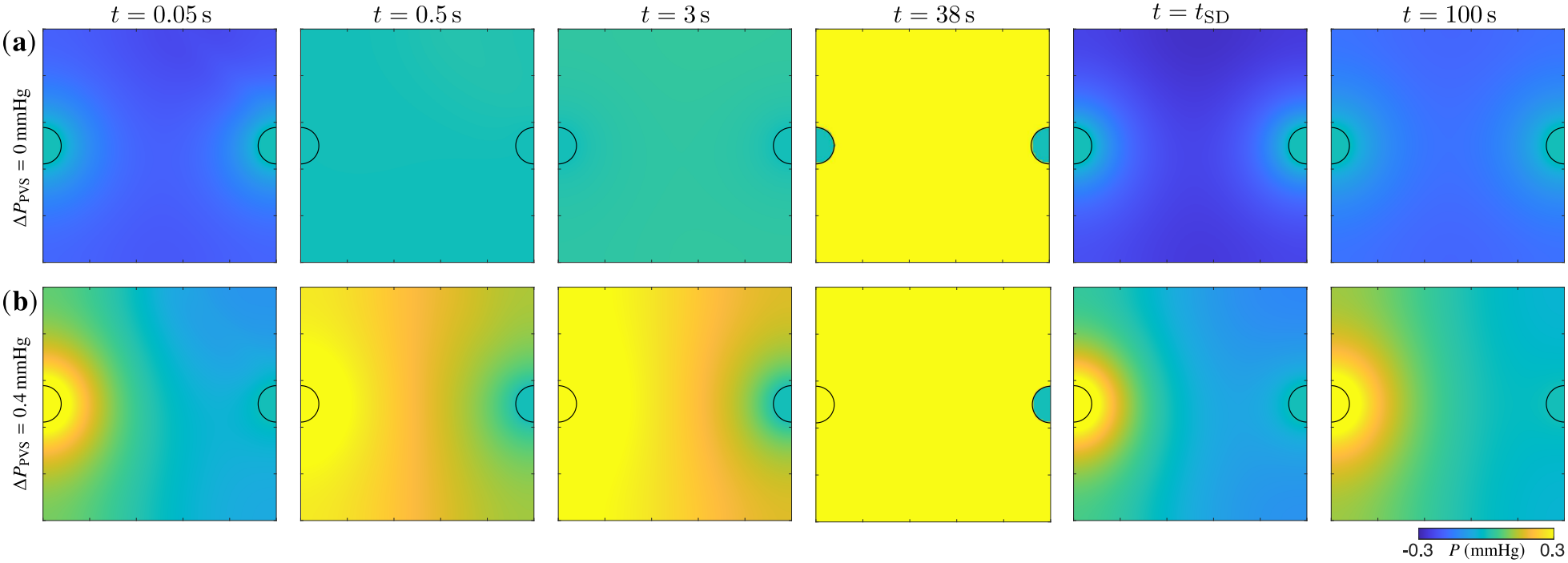
Color contours of the pressure induced by the SD wave for (a) Δ*P*_PVS_ = 0 mmHg and (b) Δ*P*_PVS_ = 0.4 mmHg. Time instances as in Fig. 7.

The color contours of the SD-induced ISF flow-magnitude with imposed streamlines are shown in Fig. 11. At the start, the reduced pressure at the initiation site rapidly draws in fluid from the surrounding PVSs, with rapid inflow from the arterial PVS for Δ*P*_PVS_ = 0.4 mmHg, and nearly symmetric inflow for Δ*P*_PVS_ = 0. As time progresses, the velocity magnitude dwindles for Δ*P*_PVS_ = 0 (Fig. 11(a)), while for the case with Δ*P*_PVS_ = 0.4 mmHg, the bulk flow driven by the PVS pressure gradient with the first-order porosity-induced contribution, drives flow from the arterial to the venous PVS. Indeed, during the recovery phase and beyond, even though the flow speed diminishes, the flow restores to the bulk-pressure-driven dipole shown in Fig. 14.

**Figure 11:**
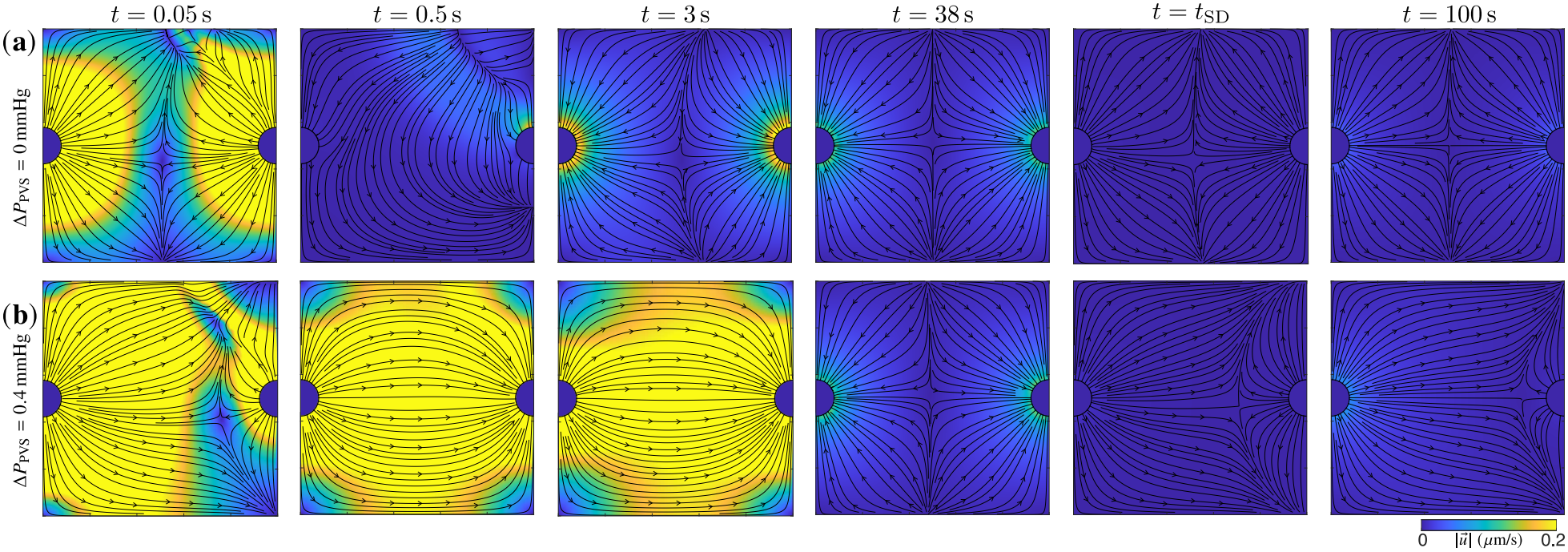
Color contours of velocity magnitude (with streamlines overlaid) induced by the SD wave for (a) Δ*P*_PVS_ = 0 mmHg and (b) Δ*P*_PVS_ = 0.4 mmHg. Time instances correspond to Fig. 7.

Figure 12(a) shows the temporal variation of the mean extracellular potassium concentration, ⟨[*K*^+^]_*e*_⟩_Ω_, which peaks at 76.32 mM at *t* = 8.8 s. Beyond *t*_*SD*_, the ⟨[*K*^+^]_*e*_⟩_Ω_ reaches a constant value of 1.44 mM that is maintained through the course of the simulation. Analogous behavior is observed for the mean membrane potential ⟨*E*_*m*_⟩_Ω_, shown in Fig. 12(b), which peaks at −10.68 mV at *t* = 9.4 s, and descends sharply at *t* = *t*_*SD*_ to a value of 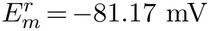. For *t* ≥ *t*_*SD*_, ⟨*E*_*m*_⟩_Ω_ increases in magnitude, reaching −78.42 mV at the end of the simulation (*t* = 100 s).

**Figure 12:**
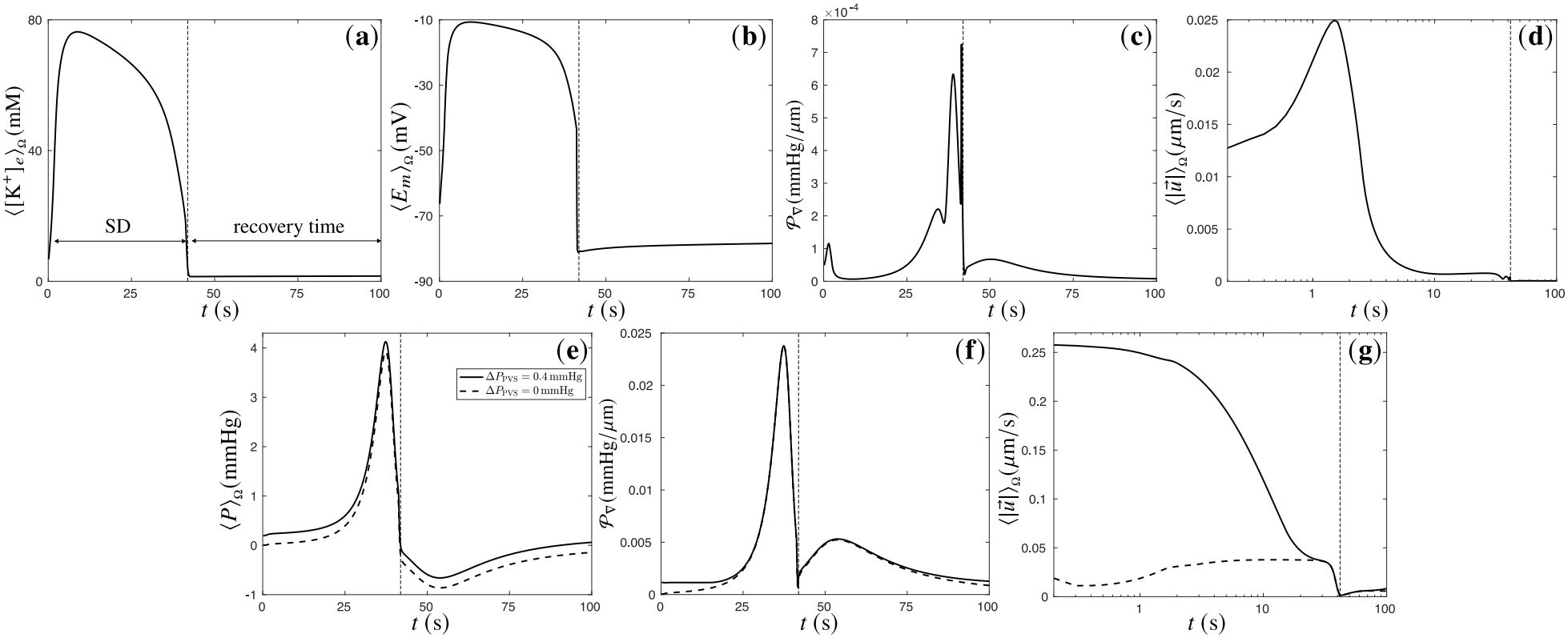
Temporal variation of ionic concentration, pressure, and velocity induced by spreading depolarization. (a) Mean extracellular potassium concentration and (b) mean membrane potential. (c) mean magnitude of pressure gradient and (d) mean velocity magnitude in a domain without PVSs. (e) mean pressure, (f) *P*_∇_, and (g) mean velocity magnitude in a domain with PVSs. Time axis is logarithmic in (d) and (g).

Figure 12(c) and (d) show the temporal variation of *P*_∇_ and 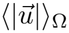 for the case without any PVSs. The mean magnitude of pressure gradient peaks at *t* ≈ 1 s, and several pronounced peaks appear as the potassium front exits the domain. These peaks originate as the pressure gradient reverses under fixed gauge ⟨*P*⟩_Ω_ = 0. The mean velocity magnitude shown in a logarithmic time axis, peaks at 0.025 *µ*m/s near the time of wave initiation at *t* = 1.5 s. Following that, the velocity drops down abruptly for *t* ≳ 8.5 s, to ≈ 7 *×* 10^−4^ *µ*m/s. For *t* ≳ *t*_*SD*_, fluid flow in the domain vanishes.

Figure 12(e-g) shows the temporal variation of the SD-induced ⟨*P*⟩_Ω_, *P*_∇_, and 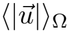, in the domain with PVSs. The mean pressure peaks at 4.13 mmHg at *t*= 37.4 s, as shown in Fig. 12(e). This increase in ISF pressure is due to the reduction in porosity, which squeezes the fluid space. Indeed, this is related to increased intracranial pressure during cytotoxic brain edema or cell-swelling [40]. During recovery, the pressure drops and becomes negative, reducing to −0.66 mmHg at *t* = 53.9 s.

The pressure then slowly recovers back to the value set by the PVS boundaries at long times.

The mean magnitude of pressure gradient increases in the course of the wave propagation, as the rising domain pressure creates a sharp gradient along the venous PVS, set at zero-pressure (Fig. 12(f)). During recovery, as the domain pressure becomes negative, it produces reverse pressure gradients from the PVSs, set at constant pressure. The mean velocity magnitude (Figure 12(g)) is set primarily by the PVSs but briefly vanishes when the SD-induced gradient cancels the imposed one near *t*_*SD*_.

We now analyze the volumetric flux through the PVS boundaries in Fig. 13. For *t* ≲ 15.8 s, Δ*P*_PVS_ = 0.4 mmHg yields substantial influx through the arterial PVS, *q*_*a*_ *<* 0 (red solid curve), with |*q*_*a*_|_*max*_ = 2.9 *×* 10^−3^ *µ*L min^−1^ mm^−1^, which is 1.2 times the arterial influx driven solely by perivascular pressure difference (dashed-dotted line). This maximum occurs at initiation, *t* = 0. As the domain pressure increases beyond the arterial PVS pressure, for 15.8 s ≤ *t* ≲ *t*_*SD*_, fluid effluxes through the arterial PVS boundary *q*_*a*_ *>* 0, reaching a maximum positive value of *q*_*a*_|_*max*_ = 5.95 *×* 10^−4^ *µ*L min^−1^ mm^−1^. For *t* ≳ *t*_*SD*_, through the end of the simulation, *q*_*a*_ ≈ 0, and is expected to recover to the resting value of PVS pressure difference-driven influx, *q*_*a*_ = −2.40 *×* 10^−3^ *µ*L min^−1^ mm^−1^, at long times.

**Figure 13:**
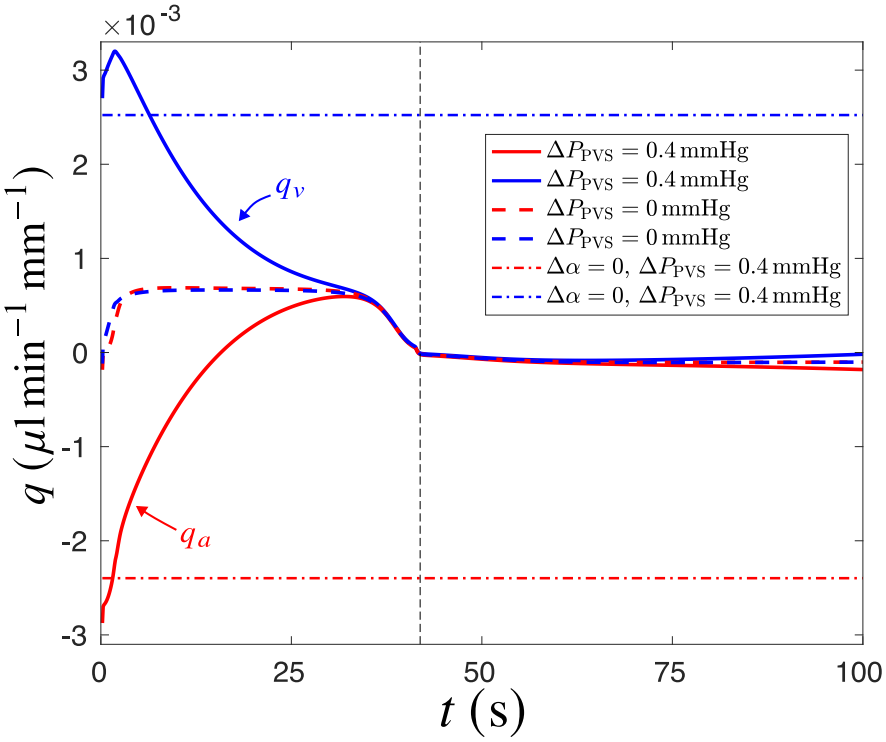
Temporal variation of the volumetric flux through the PVSs driven by the SD wave.

**Figure 14:**
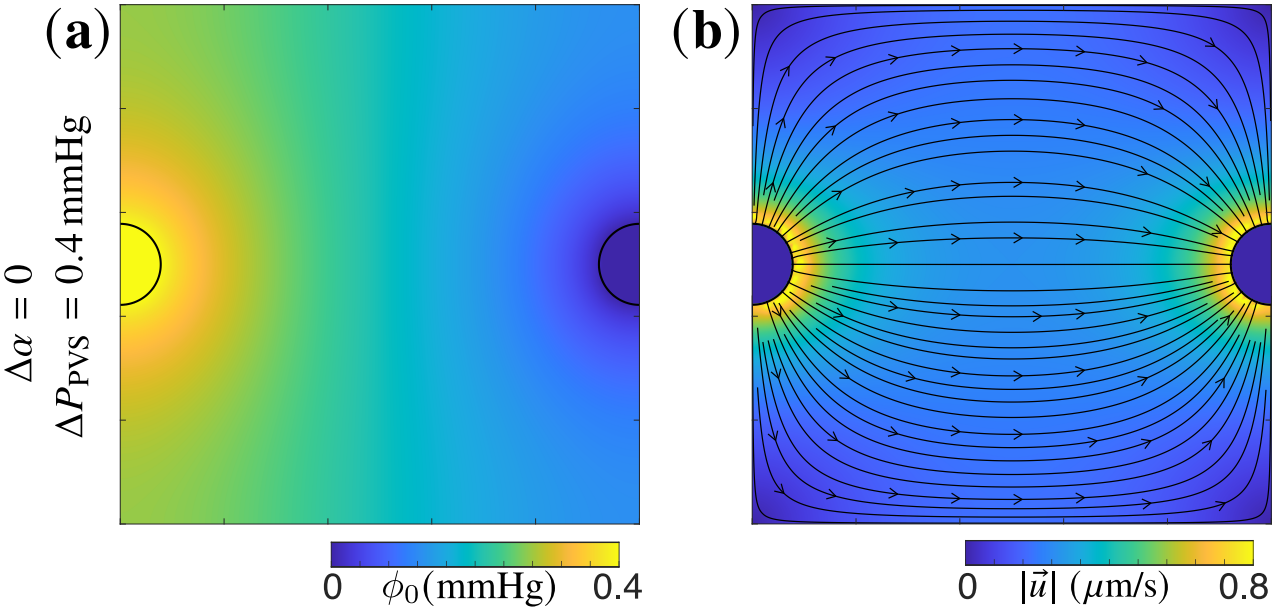
The pressure and velocity fields induced by Δ*P*_PVS_ = 0.4 mmHg without porosity variations. The pressure field equals the leading-order term *ϕ*_0_ in Eq. 60. Color contours of the (a) pressure field and (b) velocity magnitude with streamlines overlaid.

The SD-induced porosity variation also yields substantial flux through the venous PVS boundary. For *t* ≲ *t*_*SD*_, fluid effluxes through the venous PVS, with a maximum magnitude of *q*_*v*_ = 3.2 *×* 10^−3^ *µ*L min^−1^ mm^−1^ after the initiation at *t* = 2 s. For *t* ≳ *t*_*SD*_, *q*_*v*_ ≈ 0, and slowly recovers to the bulk pressure driven value at Δ*P*_PVS_ = 0.4 mmHg, of *q*_*v*_ = 2.52 *×* 10^−3^ *µ*L min^−1^ mm^−1^. For Δ*P*_PVS_ = 0, efflux through both the PVSs *q >* 0 for *t* ≲ *t*_*SD*_ is driven by the large pressure build up inside the domain driving fluid out of the zero-pressure PVS boundaries, as shown by the dotted curves.

## 4 Discussion and Conclusion

We have developed and numerically simulated a model of ISF flow driven by spatiotemporal variations in the porosity of the brain cortex. The domain represents an idealized 2D sheet of the brain cortex that is subjected to porosity variations from both a prescribed porosity variation, modeling a physiological brain wave, and a simulated spreading depolarization wave. We have presented a careful study that builds in complexity from developing the volume averaged framework, to quantifying the fluid dynamics in a domain without PVS, and a domain with PVSs at pressure differences of Δ*P*_PVS_ = 0.4 mmHg and Δ*P*_PVS_ = 0. Our intention was to probe the interaction between ionic wave and ISF flow at a level of quantitative detail that is not accessible experimentally.

We summarize our findings in Table 1. With a sinusoidal porosity wave, a 10% porosity variation drives mean flow of 0.38 *µ*m/s which rises to 0.7 *µ*m/s with PVSs at Δ*P*_PVS_ = 0.4 mmHg. Prescribed porosity variations result in an eight-fold increase in the peak volumetric fluxes through PVSs compared to that driven by Δ*P*_PVS_ alone. The response is substantially different during spreading depolarization. At the wave-initiation, mean porosity increases by ∼ 4%, and then falls sharply to 0.03. Although porosity change is larger for SD, it is far slower, and hence drives weaker flow, which scales as *∂*_*t*_*α* (Eq. (4)). In the absence of PVSs, the wave drives ISF in the direction of propagation with mean velocity reaching a peak of ≈ 0.025 *µ*m/s at *t* = 1.5 s and a peak pressure difference of Δ*P*_*max*_ = 1.36 mmHg. For Δ*P*_PVS_ = 0.4 mmHg, SD produces peak pressure difference of Δ*P*_*max*_ = 15.33 mmHg, and induces peak volumetric fluxes that are 1.3 times the value driven by Δ*P*_PVS_ alone. It also modulates the PVS-driven fluid velocity, which vanishes at *t*_*SD*_ due to the SD-induced pressure gradients canceling the imposed one. SD further induces a surge in the mean pressure, reaching over 4 mmHg, which is related to brain edema-related increase in pressure and conditions like hydrocephalus and migraine [61].

**Table 1:**
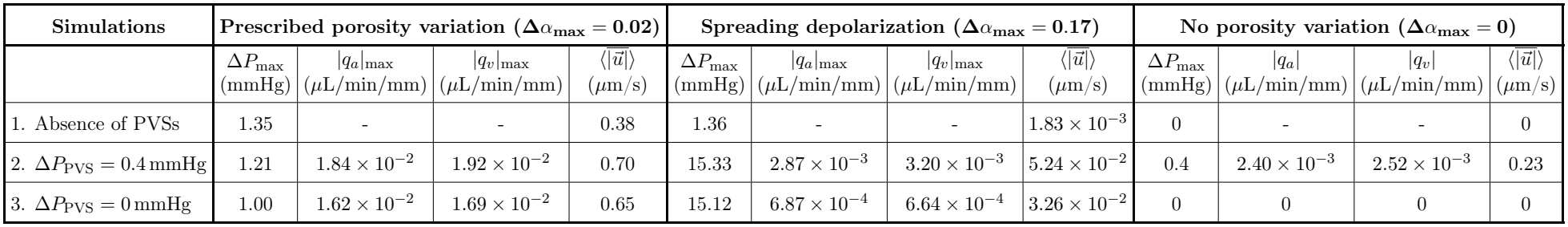
Peak pressure differences, volumetric fluxes through PVSs, and mean velocities for the different cases simulated. Δ*α*_*max*_ =|*α*−*α*_0_|_*max*_, with *α*_0_ = 0.2.

| **Simulations** | **Prescribed porosity variation (Δα<sub>max</sub> = 0.02)** |  |  |  | **Spreading depolarization (Δα<sub>max</sub> = 0.17)** |  |  |  | **No porosity variation (Δα<sub>max</sub> = 0)** |  |  |  |
|---|---|---|---|---|---|---|---|---|---|---|---|---|
|  | ΔP<sub>max</sub><br>(mmHg) | \|q<sub>a</sub>\|<sub>max</sub><br>(µL/min/mm) | \|q<sub>v</sub>\|<sub>max</sub><br>(µL/min/mm) | $\langle\overline{\|\vec{u}\|}\rangle$<br>(µm/s) | ΔP<sub>max</sub><br>(mmHg) | \|q<sub>a</sub>\|<sub>max</sub><br>(µL/min/mm) | \|q<sub>v</sub>\|<sub>max</sub><br>(µL/min/mm) | $\langle\overline{\|\vec{u}\|}\rangle$<br>(µm/s) | ΔP<sub>max</sub><br>(mmHg) | \|q<sub>a</sub>\|<br>(µL/min/mm) | \|q<sub>v</sub>\|<br>(µL/min/mm) | $\langle\overline{\|\vec{u}\|}\rangle$<br>(µm/s) |
| 1. Absence of PVSs | 1.35 | - | - | 0.38 | 1.36 | - | - | 1.83 × 10<sup>−3</sup> | 0 | - | - | 0 |
| 2. ΔP<sub>PVS</sub> = 0.4 mmHg | 1.21 | 1.84 × 10<sup>−2</sup> | 1.92 × 10<sup>−2</sup> | 0.70 | 15.33 | 2.87 × 10<sup>−3</sup> | 3.20 × 10<sup>−3</sup> | 5.24 × 10<sup>−2</sup> | 0.4 | 2.40 × 10<sup>−3</sup> | 2.52 × 10<sup>−3</sup> | 0.23 |
| 3. ΔP<sub>PVS</sub> = 0 mmHg | 1.00 | 1.62 × 10<sup>−2</sup> | 1.69 × 10<sup>−2</sup> | 0.65 | 15.12 | 6.87 × 10<sup>−4</sup> | 6.64 × 10<sup>−4</sup> | 3.26 × 10<sup>−2</sup> | 0 | 0 | 0 | 0 |

To our knowledge this is the first attempt at coupling the brain electrophysiological activities with ISF flow in the porous brain tissue using a volume-averaged formulation. A physical understanding of these contributions to the fluid dynamics will be useful on several levels, and the features identified here will guide the theoretical description of fluid dynamics in soft, absorbent porous media. Our results suggest that it may be possible to control ISF dynamics by modulating the properties of the ionic depolarization wave, for instance by externally applied neural stimulation.

## Data Availability Statement

The code used to generate the results in this paper is available at https://github.com/Drik123/SD_wave_ISF_flow.

## Authors’ Contributions

D.S. : formal analysis, investigation, methodology, validation, visualization, writing – original draft; S.M. : conceptualization, formal analysis, investigation, methodology, project administration, funding acquisition, resources, software, supervision, writing – review & editing

## Declaration of AI use

AI assistance was used for a grammar check.

## A Derivation of the governing equations of fluid flow

We use the classical volume averaging technique as derived in [56, 57] to describe the volume averaged mass and momentum conservation equation in a porous media with spatiotemporally varying porosity field. The fluid phase is denoted by *f* and the solid absorbent matrix, representing the neural and glial cells, is denoted by *s*, as indicated in Fig. 1. The volume average of any quantity in the fluid phase is

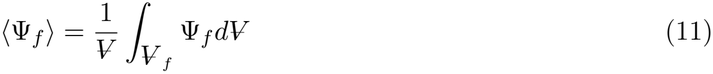

where 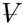 is the total averaging volume of a representative elementary volume (REV) of the solid-fluid system in the brain cortex shown in Fig. 1(b). 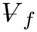 is the volume of the fluid phase (*f*) contained within the averaging volume. The REV can be considered of an arbitrary radius *r*_0_ such that *L* ≤ *r*_0_ ≤ *ℒ* [56, 57], where *L* is the characteristic length of the *f*-phase, and *ℒ* is the macroscopic length scale. For the system under consideration, we prescribe *ℒ* = 250 *µ*m, which can be considered an average distance between a penetrating arteriole and a penetrating venule in the brain cortex [22, 21, 55]. The intrinsic average, or the average over the fluid phase, of any quantity Ψ is,

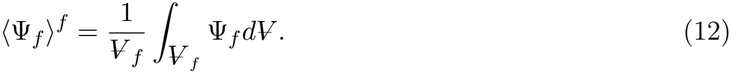

The volume fraction, or porosity, of the fluid phase *α* is defined as the ratio of the intrinsic phase average to the phase average,

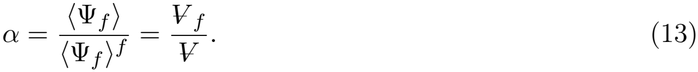

For brain tissue, the porosity is thought to be dynamic with different stages of sleep and wakefulness, with an average value of *α* ∼ 0.2 [8, 21, 22].

We will use the following averaging theorems from Refs. [56, 57, 4] in the mass and momentum conservation equations that follows this section,

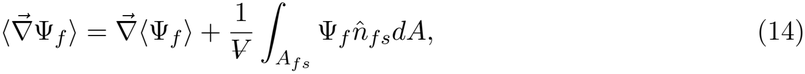

and,

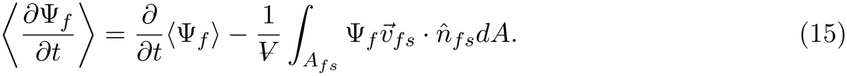

Here 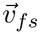 is the velocity of the interface between fluid phase and solid phase.

The continuity equation for a single-phase fluid flow through the fluid-phase of the porous medium is,

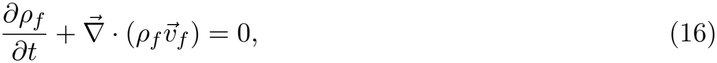

where *ρ*_*f*_ is the density and 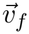 is the fluid velocity. Volume averaging the continuity equation within a representative volume (REV) leads to

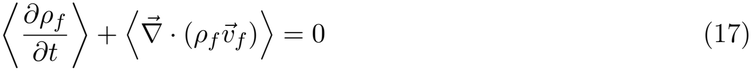

Using the averaging theorems Eqs. (14)-(15), Eq. (17) can be reduced to

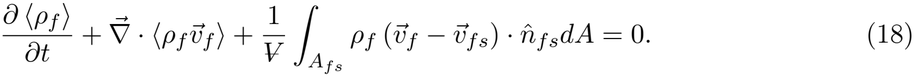

The volume average of unity is the porosity ⟨1⟩ = *α* and density is uniform, which yields:

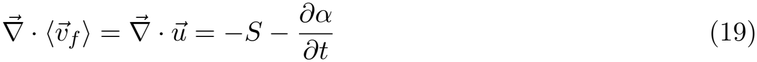

where 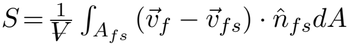 is the local imbibition rate quantifying osmotic fluid exchange between the pore space and the solid matrix due to absorption and release. For the momentum conservation, we start with the conservative form of the Navier–Stokes equation in the fluid phase, neglecting any body force,

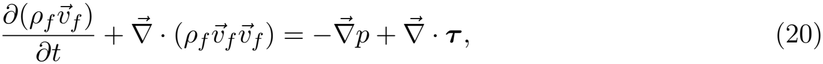

where *p* is the pressure and τ is the viscous stress tensor, such that, 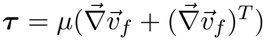, where *µ* is the coefficient of dynamic viscosity. Using the volume averaging theorems Eqs. (14)-(15), and averaging individual terms in Eq. (20), we obtain,

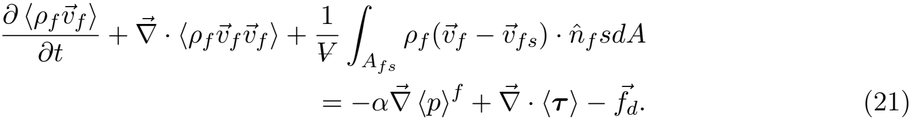

The flow through the interstitial space, over a characteristic length scale *L*, is slow and viscous with a small Reynolds number 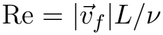 of order *O*(10^−6^), estimated with *L* ∼ 1*µ*m, *v*_*f*_ ∼ 1*µ*m*/*s, and *v* = 10^−6^m^2^*/*s. Under these conditions, inertial effects are negligible, and the transient and convective terms in the left-hand side of Eq. (21) can be neglected. The momentum conservation equation therefore reduces to,

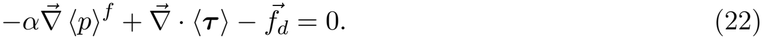

In Eq. (22), 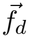 is the drag force vector obtained by combining the remnant terms from the volume averaging, such that,

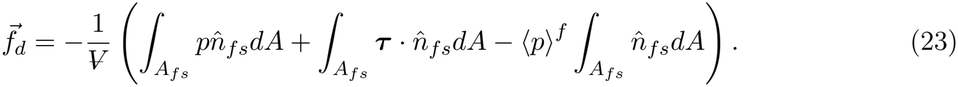

The drag can be closed following the treatment by Whitaker [56] in deriving Darcy’s law: 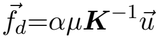, where ***K*** is the second-order permeability t ensor. Volume averaging of the shear stress term yields 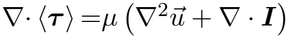 , where 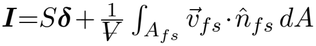, is the interfacial kinetic effects tensor [4], related to the net force acting on the interface due to the distribution of the non-zero velocity 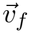 over the interface.

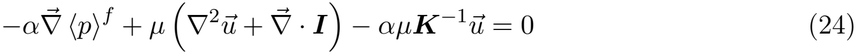

where 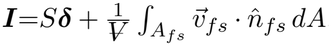, is the interfacial kinetic effects tensor [4], related to the net force acting on the interface due to the distribution of the non-zero velocity 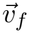 over the interface. We now scale Eq. (24), against the viscous stress scale 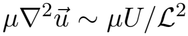, where *U* is the characteristic velocity scale. This yields a Darcy number, 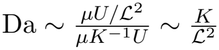, that multiplies the porous drag term. For the brain tissue, the permeability is of the order of *K* = *O*(10^−17^) m^2^ [23] and *ℒ* = *O*(10^−3^) m, which yields Da∼ 10^−11^. This implies that the viscous term will not lead to any substantial pressure difference, and we can further reduce the equation to,

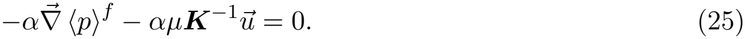

Indeed, this reduced form of the momentum equation is equivalent to the Darcy’s law with a spatiotemporally variable permeability. Writing the intrinsic average of the pressure as *P* = ⟨*p*⟩^*f*^ , and hydraulic conductivity, *k* = *K/µ*, yields the momentum equation,

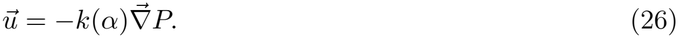

## B Numerical implementation

We use a finite difference approach to solve the governing equations with second-order accuracy in space using central differences and fourth-order accuracy in time using explicit Runge–Kutta time marching. The approach has been implemented in Fortran and has been numerically verified [42]. For the prescribed porosity field Eq. (6), a time step of 0.005 ms, and a spatial grid size of 101*×*101 points is used. For the spreading depolarization wave, we use a grid size of 26 *×* 26 to solve Eqs. (67)-(72). We then interpolate the resultant ion and porosity fields onto a 101 *×* 101 spatial grid, and use them to solve for the corresponding pressure and flow fields. In this second stage, the time step is varied between 1ms and 0.1s, depending on the temporal resolution required and the timescale of the simulated dynamics. At each time step of the wave, the pressure Poisson equation has been solved using a Gauss–Seidel iteration scheme to get the local flow field perturbation induced by the traveling wave.

The boundary condition for the pressure field is set to a Neumann boundary condition 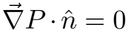 on all bounding surfacesΓ = *∂*Ω, where 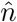 is the generalized outward pointing normal at a surface. This implies that the surfaces are impermeable and that the component of fluid velocity parallel to the boundary is nonzero, but the perpendicular component is zero. When PVSs are absent, we fix the gauge at each time step by imposing a zero mean pressure ⟨*P*⟩_Ω_ = 0. If PVSs are present, a Dirichlet condition is used in the arterial and venous PVS boundaries,Γ_*a*_ ⊂ *∂*Ω, andΓ_*v*_ ⊂ *∂*Ω, and it is not required to fix a gauge pressure. We study two scenarios: (1) a perivascular pressure difference of Δ*P*_PVS_ = 0, where arterial and venous PVSs are both set to *P* = 0 mmHg, and (2) Δ*P*_PVS_ = 0.4 mmHg, where arterial PVS is set to *P* = 0.4 mmHg and venous PVS is set to *P* = 0 mmHg. The arterioles and venules are assumed to be circles of diameter 30 *µ*m, each with an annular PVS of width 5 *µ*m surrounding the vessels [24, 21, 22]. The initial concentration of the ionic concentrations are set to the corresponding resting values. For initiating spreading depolarization, the extracellular potassium concentration at *t* = 0 is set to a localized Gaussian function, given as

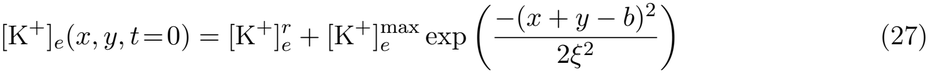

simulating a spike in potassium concentration at a localized region in space. Here, *b* = 0.04 cm, *ξ* = 5 *×* 10^−4^ cm, and the initial spike is of magnitude 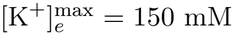. This initial condition replicates the scenario of KCl application to instigate SD waves in experiments [62]. The initial condition for the pressure field is either *P* = 0 everywhere in the domain, or when perivascular pressure gradient is imposed, the PVSs are set to their boundary values for *t <* 0.

## C. Analytical derivation of pressure field due to small change in α in the large-wavelength limit ℒ ≪ λ

We consider the governing pressure equation

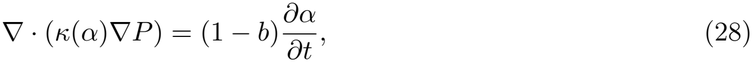

where the hydraulic conductivity is given by the Kozeny–Carman relationship Eq. (5)

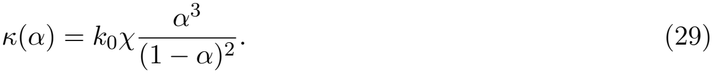

We consider the porosity to evolve as the prescribed porosity field Eq. 6 and consider a small perturbation about the constant base state *α*_0_:

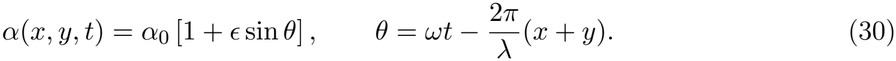

Therefore,

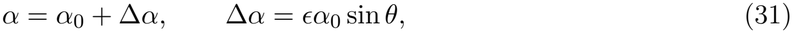

where Δ*α* is the small change and *α*_0_ is a baseline value of porosity. The time derivative of *α* is

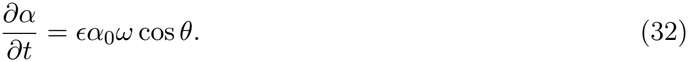

We expand *κ*(*α*) about *α*_0_, using a Taylor series:

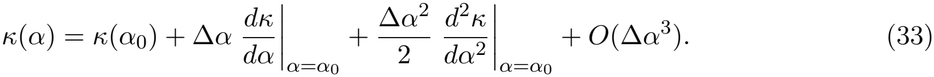

The first and second derivatives of *κ*(*α*) are given as

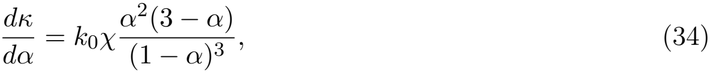

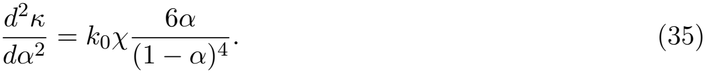

To solve for the unknown pressure field, we use a straightforward expansion in the small parameter *ϵ*:

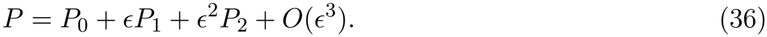

Consequently,

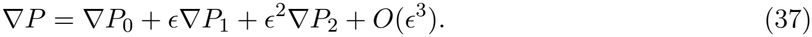

Substituting Eqs. (32), (33) and (37) in Eq. (28), with Eqs. (34) and (35) substituted in Eq. (33), we get

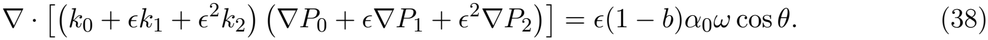

Where

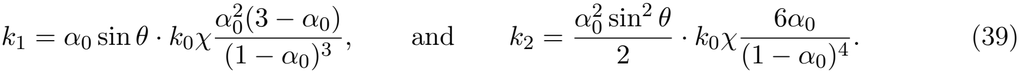

Collecting terms by order gives the leading-order equation as

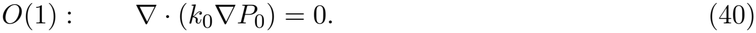

Since *κ*_0_ is constant, this reduces to

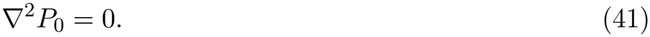

Since BCs are constant with time, this leads to

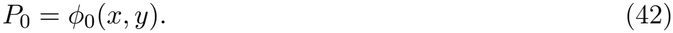

At first order of Eq. (38), we obtain

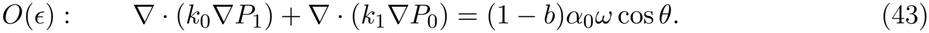

In this equation, *k*_0_ is constant, thus the first term in the left hand side can be rewritten as *k*_0_∇^2^*P*_1_. For the second term, only the sin *θ* term in *k*_1_ (from Eq. (39)) is a function of space. Thus, from Eq. (43), we get

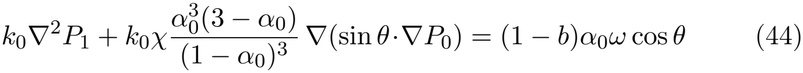

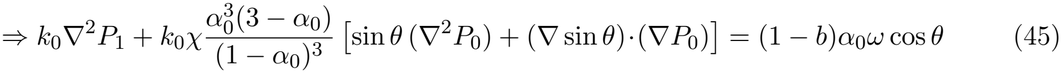

From Eq. (41), we have ∇^2^*P*_0_=0. And, assuming large wavelength (*λ* ≫ *L*), the second term in the square brackets derive as (∇ sin *θ*) = (2*π/λ ·* cos *θ*) = 0. Thus, simplifying Eq. (43), both the terms inside the square brackets equal 0, yielding,

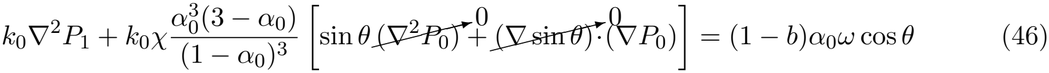

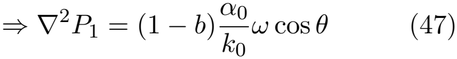

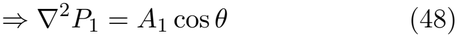

The particular solution to *P*_1_ is given as:

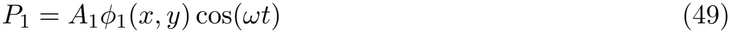

where

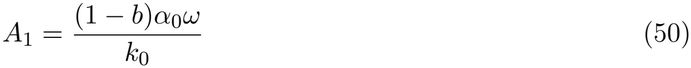

Collecting the second-order terms in Eq. (38) gives

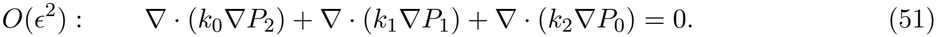

Since *k*_0_ is constant, this can also be written as

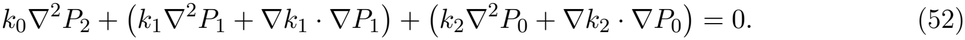

Since the leading-order pressure is assumed to be spatially constant,

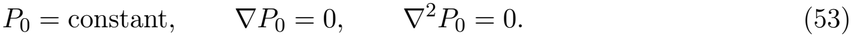

Therefore, the terms involving *P*_0_ vanish, and Eq. (52) reduces to

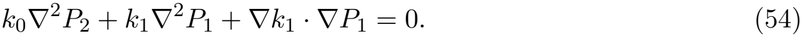

Here, the third term comprising a (∇ sin *θ*) expression also vanishes the same way as in Eq. (46). Using the first-order pressure equation, Eq. (47), the second-order equation leads to

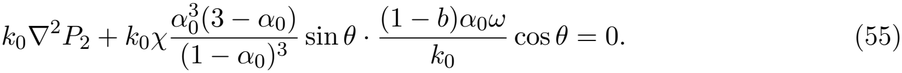

Upon simplifying this equation, we get

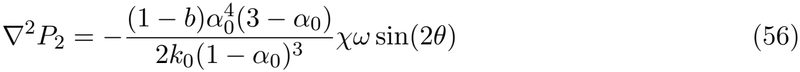

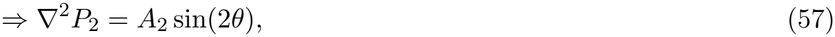

where,

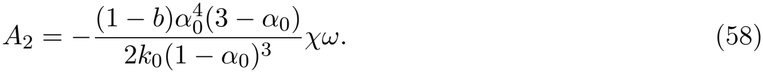

And the particular solution from Eq. (57), since *ℒ/λ* → 0, leads to

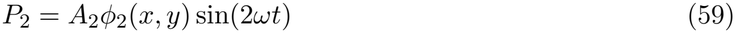

Also, *ϕ*_1_ and *ϕ*_2_ are solutions to the same equations ∇^2^*ϕ*_1_ = ∇^2^*ϕ*_2_ = 1, with the same boundary conditions (B.C.s), thus, *ϕ*_2_ = *ϕ*_1_. Thus, the solution to *P* is,

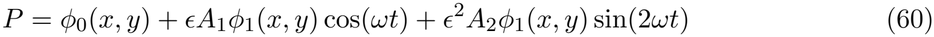

And the spatially averaged definition for *P* is:

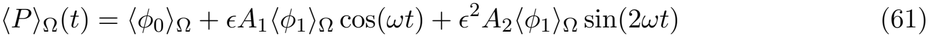

where ⟨*ϕ*_0_⟩_Ω_ and ⟨*ϕ*_1_⟩_Ω_ are constants, since *ϕ*_0_ and *ϕ*_1_ are solely functions of *x* and *y*. The spatially averaged quantities ⟨*ϕ*_0_⟩_Ω_ and ⟨*ϕ*_1_⟩_Ω_ are calculated after solving for *ϕ*_0_ and *ϕ*_1_ using the following equations

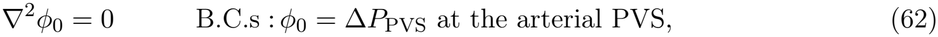

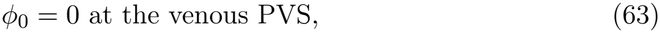

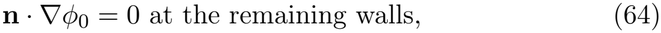

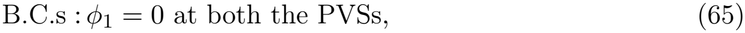

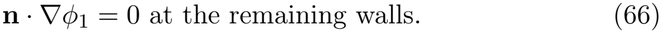

The leading order term *ϕ*_0_(*x, y*) is numerically solved for Δ*P*_PVS_ = 0.4 mmHg and is shown in Fig. 14. We now compare the numerical results we obtain by solving the full equation with prescribed porosity variation (Eq. (6)), and the large-wavelength analysis Eq. (60). The solid lines in Fig. 15 show ⟨*P*⟩_Ω_(*t*), obtained from Eq. (61), while the data points show the numerical simulation results from Sec. 33.1. While a reasonable agreement is found, the departures from the large-wavelength solution is due to the finite wavelength of *λ* = 1 mm that is considered, especially near *ωt* = *π/*2 and *ωt* = 3*π/*2.

**Figure 15:**
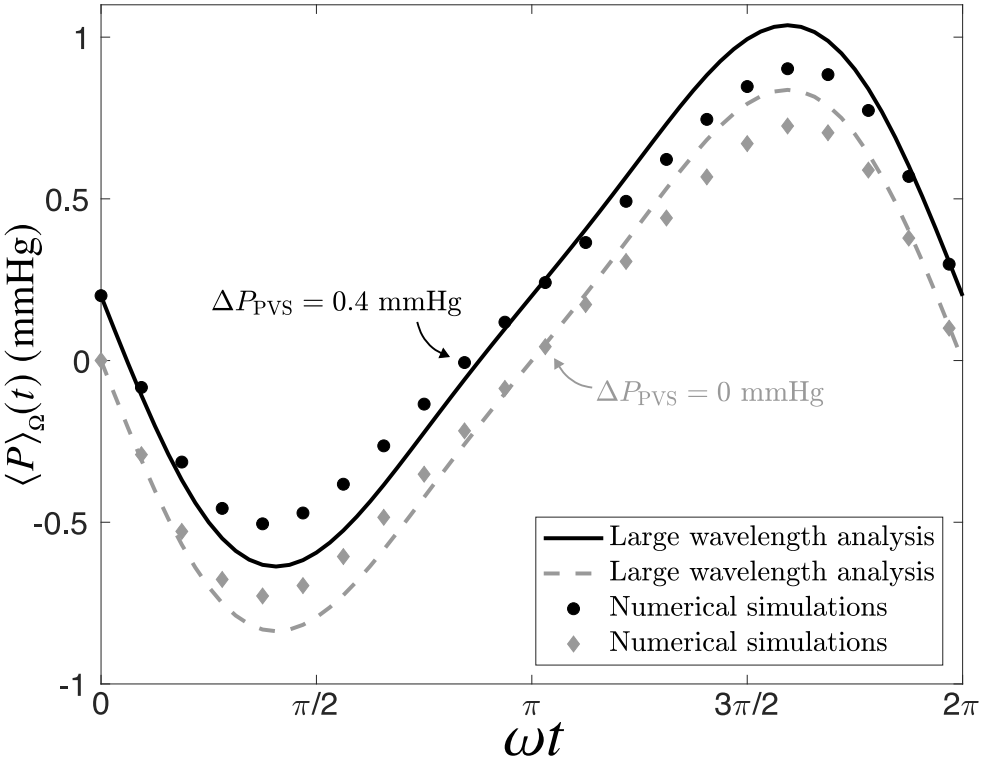
Mean pressure versus time for Δ*P* = 0.4mmHg and Δ*P* = 0, obtained from large-wavelength analysis.

## D Governing equations for ionic dynamics during spreading depolarization

Incorporating this dynamic porosity, the ionic transport can be represented by the following set of equations [42, 60, 11]:

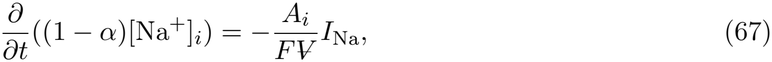

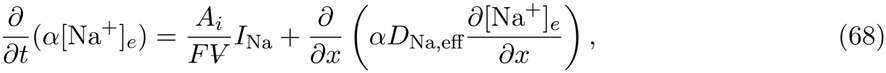

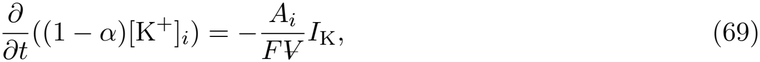

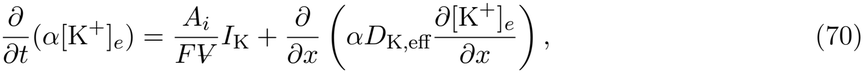

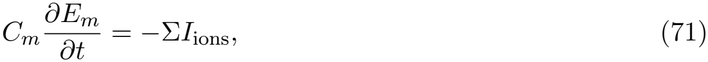

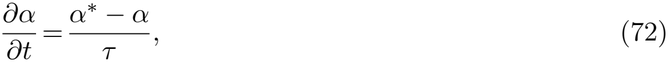

where *e* and *i* denote extra- and intracellular space, respectively, *E*_*m*_ is the cell membrane potential, *C*_*m*_ is the membrane capacitance, *A*_*i*_ is the surface area of the cells, *F* is Faraday’s constant, *D*_*ion,eff*_ = *D*_*ion*_/Λ^2^ is the effective diffusion coefficient of ions in the porous media, with Λ = 2.04 being the tortuosity of the brain interstitium [7]. S*I*_*ions*_ =*I*_*Na*_+*I*_*K*_+*I*_*leak*_ in Eq. (71) represents the total cross-membrane ionic currents due to the ionic movements of potassium and sodium ions across the membrane between the cells and extracellular space. *I*_*leak*_ is the general leak current, while *I*_*Na*_ and *I*_*K*_ represent the sodium and potassium currents, respectively, comprising the individual active, leak and pump currents [42, 34, 35]. The sodium and potassium currents consist of active currents which are modeled by GHK equations and passive currents by HH equations. These currents are modulated by the gating variables in the cell membrane, which evolve in time. We direct the reader to Ref. [42] for these details. The resulting framework, yields a 14-component reaction-diffusion system which we solve in Fortran.

Equation (72) defines the rate of change of porosity, treated as a first-order process with a time constant of *τ* = 250 ms [11], and is motivated by electrophysiological models of cell swelling during seizures, anoxia and spreading depolarization [11, 10]. With the assumptions of reference brain porosity *α*_0_, the total sum of extra and intracellular volume being constant at 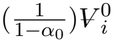, where 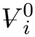 is the initial neuronal volume, and the extracellular volume maximally shrinking to *η* times the initial cellular volume at spreading depression [11], the expected porosity *α*^*^ due to the difference between extracellular and intracellular osmolarities *π*_*e*_ and *π*_*i*_, given as

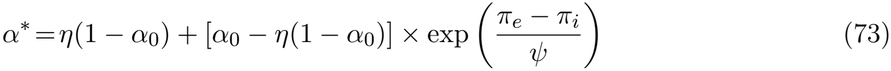

where *ψ* = 20 is a phenomenological choice [35], and the intra- and extracellular osmolarities are governed by the ionic salt concentrations of sodium and potassium, defined as

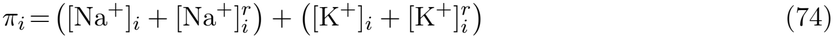

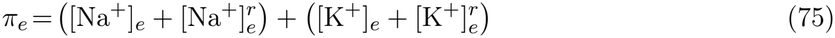

where the resting values of the ionic concentrations, are given as, 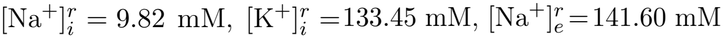 and 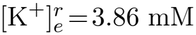 [34, 60]. It is important to note that the ionic evolution does not depend on the ISF flow since ions are small, and diffusion is orders of magnitude faster than advection [7, 42]

